# Essential Yet Dispensable: The Role of CINNAMATE 4-HYDROXYLASE in Rice Cell Wall Lignification

**DOI:** 10.1101/2024.10.08.617307

**Authors:** Supatmi, Lydia Pui Ying Lam, Senri Yamamoto, Osama A. Afifi, Pingping Ji, Yuriko Osakabe, Keishi Osakabe, Toshiaki Umezawa, Yuki Tobimatsu

## Abstract

A comprehensive understanding of the intricate lignin biosynthesis in grasses could contribute to enhancing our ability to utilize grass biomass. CINNAMATE 4-HYDROXYLASE (C4H), in conjunction with PHENYLALANINE AMMONIA-LYASE (PAL), initiates the entry of phenylalanine into the cinnamate/monolignol pathway, leading to the production of diverse phenylpropanoids, including lignin monomers. Despite extensive research on C4H in eudicots, genetic studies of C4H in grasses remain considerably limited. Notably, the role of C4H in the presence of PHENYLALANINE/TYROSINE AMMONIA-LYASE (PTAL), a grass-specific ammonia-lyase that can bypass the conserved PAL-C4H pathway by recruiting tyrosine into the cinnamate/monolignol pathway, remains unclear. To address this gap, a set of genome-edited rice mutants harboring knockout mutations in rice *C4H* genes were generated and subjected to the analysis of growth phenotype and cell wall chemotype, alongside isotopic feeding and chemical inhibitor assays to test the contributions of the PAL-C4H and PTAL pathways. The phenotype and chemotype characterizations of *C4H*-knockout rice mutants demonstrated that class I (OsC4H1/CYP73A38) and class II (OsC4H2a/CYP73A39 and OsC4H2b/CYP73A40) C4Hs cooperatively contribute to lignin biosynthesis in rice. Nevertheless, the impacts of *C4H*-deficiency on plant development and lignin formation in rice appeared to be less prominent compared to those reported in eudicots. The ^13^C-labeled phenylalanine and tyrosine feeding experiments demonstrated that even with the phenylalanine-derived PAL-C4H pathway completely blocked, the *C4H*-knockout rice can still produce significant levels of lignin and maintain sound cell walls by utilizing the tyrosine-derived PTAL pathway. Overall, this study demonstrates the essential but dispensable role of C4H in grass cell wall lignification.

## Introduction

Lignin is a phenylpropanoid polymer produced in the secondary cell walls of vascular plants, typically accounting for 15–30% of lignocellulosic biomass. While lignin is a vital component in plant cell walls, providing essential mechanical strength, water conductivity, and resistance to pathogens, it has long been considered a recalcitrant component in polysaccharide-oriented biomass utilizations such as wood pulping and fermentable sugar productions (Boerjan *et al*., 2003; Umezawa, 2010; Bonawitz and Chapple, 2013; Halpin, 2019; Mahon and Mansfield, 2019). More recently, however, lignin has been increasingly viewed also as a viable source for the production of energy-rich fuels as well as valuable aromatic chemicals in the light of the biorefinery concept (Rinaldi et al., 2016; Umezawa, 2018; Umezawa et al., 2020; Abu-Omar et al., 2021; Wang et al., 2022; Umezawa, 2024).

Diverse phenylpropanoids derived from the cinnamate/monolignol pathway serve as lignin monomers that combine to form complex lignin polymers via oxidative radical coupling in the cell walls (Ralph et al., 2004; Tobimatsu and Schuetz, 2019). The formulation of lignin monomers, which is the primary determinant of the structure and properties of lignin polymers, varies among the classes of vascular plants (Ralph et al., 2019). In general, lignins produced by most ferns, gymnosperms, and eudicots are derived mainly from monolignols, namely coniferyl, sinapyl, and *p*-coumaryl alcohol, which give rise to guaiacyl (G), syringyl (S), and *p*-hydroxyphenyl (H) units, respectively, in the final lignin polymers (**Figure 1**). On the other hand, lignins produced by monocotyledonous grasses additionally use monolignol *p*-hydroxycinnamate conjugates, mainly γ-*p*-coumaroylated monolignols (coniferyl and sinapyl *p*-coumarate) (**Figure 1**) (Ralph et al., 1994; Ralph, 2010; Karlen et al., 2018; Chandrakanth et al. 2023). Furthermore, grass lignin uniquely incorporates tricin, a flavonoid, which copolymerizes with monolignols and γ-p-coumaroylated monolignols to be integrated into the lignin polymer structures. (**Figure 1**) (del Río et al., 2012; Lan et al., 2015; del Río et al., 2020; Lam et al., 2021; Lam et al., 2023). Because of the considerable interest in applying molecular breeding and bioengineering approaches to control grass lignin biosynthesis for agro-industrial applications of grass biomass, the regulatory mechanisms of the cinnamate/monolignol pathway that coordinate the production of diverse lignin monomers in grasses have been the focus of the related research (Dixon and Barros, 2019; Umezawa et al., 2020; Lam et al., 2021; Chandrakanth et al. 2023; Umezawa, 2024).

**Figure 1.**
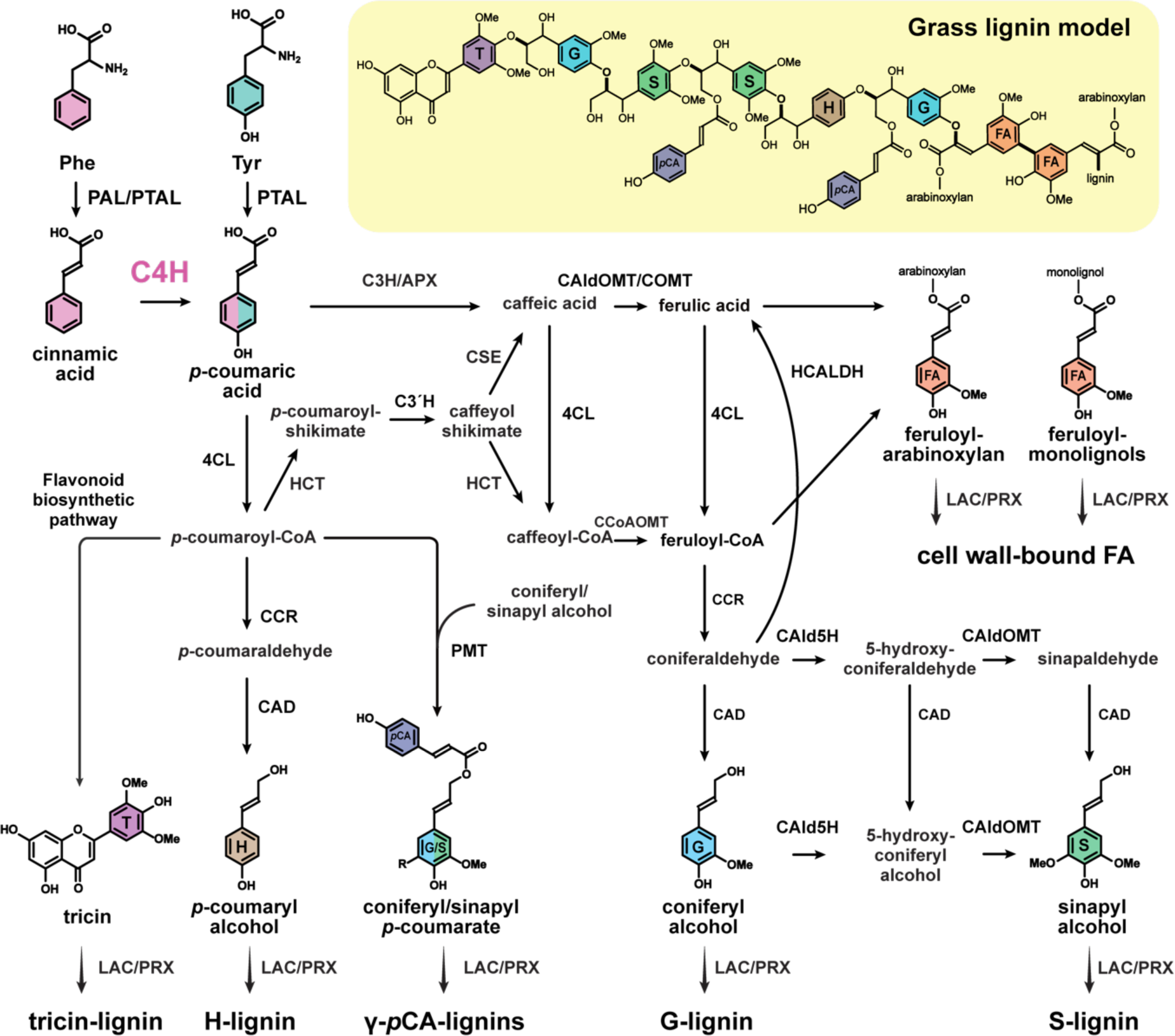
Proposed biosynthetic pathways to lignin monomers in grasses. PAL, phenylalanine ammonia-lyase; PTAL, phenylalanine/tyrosine ammonia-lyase; C4H, cinnamate 4-hydroxylase; 4CL, 4-coumarate:CoA ligase; C3H/APX, 4-coumarate 3-hydroxylase/ascorbate peroxidase; C3’H, p-coumaroyl ester 3-hydroxylase; HCT, *p*-hydroxycinnamoyl-CoA:quinate/shikimate transferase; CSE, caffeoyl shikimate esterase; CCoAOMT, caffeoyl-CoA *O*-methyltransferase; CCR, cinnamoyl-CoA reductase; HCALDH, hydroxycinnamaldehyde dehydrogenase; CAld5H, coniferaldehyde 5-hydroxylase; CAD, cinnamyl alcohol dehydrogenase; CAldOMT/COMT, 5-hydroxyconiferaldehyde *O*-methyltransferase/caffeic acid *O*-methyltransferase; PMT, *p*-coumaroyl-CoA:monolignol transferase; LAC, laccase; PRX, peroxidase; H, *p*-hydroxyphenyl; G, guaiacyl; S, syringyl; T, tricin; *p*CA, *p*-coumarate; FA, ferulate.

In the proposed cinnamate/monolignol pathway, CINNAMATE 4-HYDROXYLASE (C4H), a cytochrome P450 (CYP) enzyme belonging to the CYP73A family, catalyzes the *para*-hydroxylation of cinnamic acid into *p*-coumaric acid (Fahrendorf and Dixon 1993; Mizutani et al., 1993; Teutsch et al., 1993; Werck-Reichhart et al., 1993). Alongside PHENYLALANINE AMMONIA-LYASE (PAL), C4H initiates the entry of phenylalanine (Phe) into the cinnamate/monolignol pathway dedicated for the biosynthesis of diverse phenylpropanoids, including cell wall polymers, such as lignin and suberin, as well as diverse soluble phenolics, such as flavonoids and stilbenoids (**Figure 1**). The activities of C4H therefore can affect the entire phenylpropanoid biosynthetic pathways and thereby the production of the downstream phenylpropanoids, contributing to many aspects of plant development and physiology. C4H may interact with other CYP enzymes in the cinnamate/monolignol pathway (Chen et al., 2011) and scaffold proteins (Gou et al., 2018; Zhao et al., 2023) to form enzyme complexes or metabolons, potentially influencing metabolic flux through the pathway’s branches. Due to its important roles in lignin and other phenylpropanoid biosynthesis, C4H has been extensively studied across various plant species.

A number of genetic studies of C4H homologs in eudicots, including Arabidopsis (*Arabidopsis thaliana*) (Schilmiller et al., 2009; El Houari et al., 2021; Kim et al., 2021), tobacco (*Nicotiana* spp.) (Sewalt et al., 1997; Blee et al., 2001), poplar (*Populus* spp.) (Bjurhager et al., 2010; Kim et al., 2020), alfalfa (*Medicago sativa*) (Reddy et al., 2005), and eucalyptus (*Eucalyptus* spp.) (Sykes et al., 2015; Ziebell et al., 2016) and so on, have demonstrated the essential role of C4H in lignin biosynthesis in these plants. Typically, down-regulation of *C4H* genes in eudicots results in substantial reductions in lignin contents in the cell walls, often accompanied by impaired vasculature and plant growth depending on the degree of gene suppression. In Arabidopsis, a *C4H*-knockout T-DNA mutant showed seedling lethality and early arrest of leaf expansion (Schilmiller et al., 2009). The *reduced epidermal fluorescence 3* (*ref3*) Arabidopsis mutants with leaky expression of *C4H* showed substantially reduced lignin levels, down to ∼20% of wild-type levels, along with severe growth phenotypes, such as abnormal stem and leaf development, collapsed vasculature, male sterility, and dwarfism (Schilmiller et al., 2009; El Houari et al., 2021; Kim et al., 2021; El Houari et al., 2021). These data strongly suggest that C4H is indispensable for lignin biosynthesis and sound plant development in eudicots.

Despite the extensive research on C4H in eudicots, genetic and biochemical studies of C4H in monocotyledonous grasses remain considerably limited. C4H proteins from the model grass *Brachypodium distachyon* displayed typical C4H enzymatic activity *in vitro* and could complement the growth defect of a *C4H*-deficient Arabidopsis mutant (Renaut et al., 2017; Knosp et al., 2024). Nevertheless, a lack of information on *C4H*-downregulated grass mutants hinders our understanding of C4H’s specific contribution to grass lignin biosynthesis. As aforementioned, unlike eudicots, grasses employ a more complex cinnamate/monolignol pathway to generate both common monolignols and lineage-specific lignin monomers such as tricin (a flavonoid) and *p*-hydroxycinnamate conjugates for cell wall lignification (**Figure 1**). Notably, unlike eudicots (and other most vascular plants) that rely on the conserved Phe-derived PAL-C4H pathway for lignin biosynthesis, grasses possess a unique parallel pathway, which bypasses the need for the Phe-derived PAL-C4H pathway by employing the PHENYLALANINE/TYROSINE AMMONIA-LYASE (PTAL) to produce *p*-coumaric acid from tyrosine (Tyr) (Barros et al., 2016; Barros and Dixon, 2020; El-Azaz et al., 2023) (**Figure 1**). The interplay between these dual lignin pathways, i.e., the Phe-derived PAL-C4H pathway and the Tyr-derived PTAL pathway, in coordinating the production of diverse grass lignin monomers remains elusive.

This study investigates the role of C4H in rice (*Oryza sativa*), a model grass species. Using CRISPR-Cas9-mediated targeted mutagenesis, we generated rice mutants with single and multiple knockouts of the three major *C4H* genes. To test the involvements of the rice C4Hs in lignin biosynthesis and cell wall development, we then analyzed the growth phenotypes of the *C4H*-knockout rice mutants and performed in-depth cell wall structural analyses using chemical and two-dimensional (2D) nuclear magnetic resonance (NMR) techniques. To elucidate the contributions of the dual PAL-C4H and PTAL pathways, we employed isotopic feeding assays using ^13^C-labeled Phe and Tyr substrates on both *C4H*-knockout and wild-type rice plants. Additionally, chemical inhibitor assays utilizing C4H and PAL/PTAL inhibitors were conducted.

## Results

### Sequence and gene expression analysis of rice C4Hs

We initiated this study by examining the protein sequences and gene expression of rice C4H (CYP73A) family members. The rice genome encodes four putative C4H proteins: OsCYP73A35p, CYP73A38 (hereafter referred to as OsC4H1), CYP73A39 (hereafter referred to as OsC4H2a), and CYP73A40 (hereafter referred to as OsC4H2b). Phylogenetic reconstruction classified OsCYP73A35p and OsC4H1 into the grass class I C4H proteins and OsC4H2a and OsC4H2b into the grass class II C4H proteins along with other putative C4H proteins in grasses (**Figure 2A; Supplemental Table S1**). These rice C4H proteins share over 58% sequence identity with each other (**Supplemental Table S2**). The two class II C4H, OsC4H2a and OsC4H2b share approximately 99% amino acid sequence identity, suggesting that they are likely products of a recent gene duplication event. Multiple alignment indicated that OsC4H1, OsC4H2a, and OsC4H2b possess conserved cytochrome P450 sequences, including proline cluster, oxygen binding domain, and heme binding domain, essential for C4H activity (Chapple, 1998) (**Supplemental Figure S1**). In contrast, CYP73A35p possesses a truncation of approximately 20 amino acid residues in its functional domain and lacks the oxygen-binding region (**Supplemental Figure S1**), suggesting that it is a non-functional C4H protein encoded by pseudogene.

**Figure 2.**
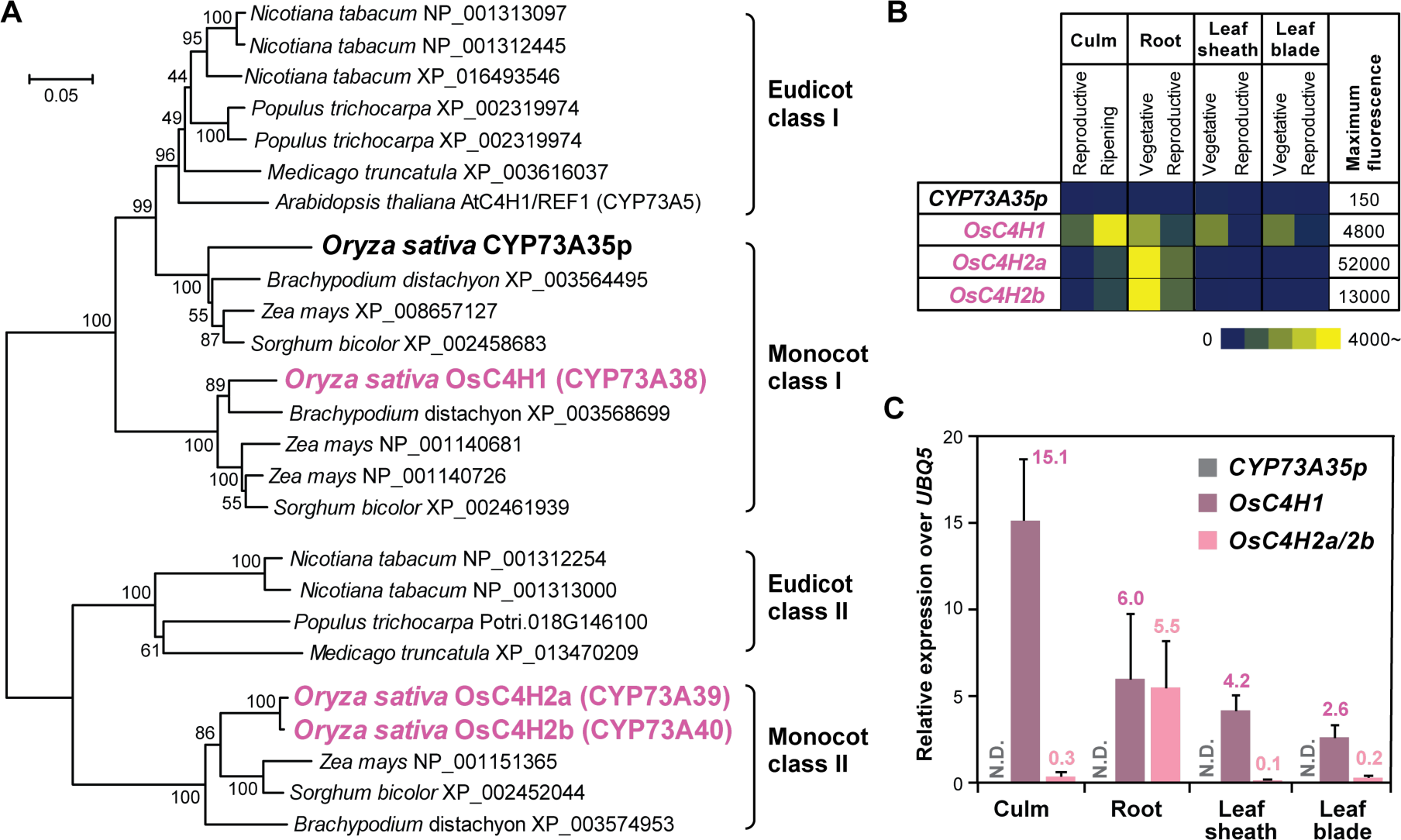
Phylogeny and gene expression analysis of rice C4H members. **A**) Phylogenetic tree of C4H (CYP73A) proteins. The unrooted phylogenetic tree of C4H proteins from rice and other major grass and eudicot species were built by the maximum-likelihood method with bootstrap values after 1,000 tests. Scale bar denotes 0.05 substitutions per site. Gene locus and protein accession numbers are listed in **Supplemental Table S1**. **B**) *In silico* relative expression of rice *C4H* genes retrieved from the public DNA microarray database, RiceXPro (Sato et al., 2013). **C**) Relative expression of three *C4H* genes in the wild-type rice at the heading stage as determined by RT-qPCR. A ubiquitin gene (*OsUBQ5*) was used as an internal control. Values are mean ± standard deviation of biological replicates (*n* = 3).

Next, we examined the spatio-temporal expression pattern of rice *C4H* genes using publicly available gene expression data from the RiceXpro gene expression database (Sato et al., 2010). Among the two class I *C4H* genes, *OsC4H1* exhibited relatively high expression levels in major vegetative organs (culm, root, leaf sheath, and leaf blade), with the highest expression level in the culm. In contrast, the expression levels of *CYP73A35p* in those organs were quite low (**Figure 2B**). The two class II *C4H* genes, i.e., *OsC4H2a* and *OsC4H2b*, also showed high expression in the root, with considerably lower expression levels in the aerial organs (culm, leaf sheath, and leaf blade) (**Figure 2B**). Gene expression analysis using reverse transcription-quantitative PCR (RT-qPCR) verified these expression patterns of rice *C4H* genes (**Figure 2C**). Expression of *OsC4H1* was detected in the major rice vegetative organs (culm, root, leaf sheath, and leaf blade) examined, whereas expression of *OsC4H2a* and *OsC4H2b* was more specifically detected in the root. Consistent with the *in silico* gene expression data, expression of *CYP73A35p* was not detected in any of the examined rice organs. Based on these protein sequence and gene expression data, we excluded *CYP73A35p* from further functional characterization.

### Generation of *C4H*-knockout rice mutants via CRISPR-Cas9-mediated mutagenesis

To investigate the *in-planta* functions of rice C4H members, we generated genome-edited rice mutant lines deficient in *OsC4H1*, *OsC4H2a*, and *OsC4H2b*. Single-guide RNAs (sgRNAs) that target the conserved C4H active site regions (Chapple, 1998) were designed for each of the *C4H* sequences using the CRISPR-P 2.0 program (Liu et al., 2017) (**Supplemental Table S3**). Due to the high similarity between the *OsC4H2a* and *OsC4H2b* sequences, we designed sgRNAs that target their common sequence to edit both genes simultaneously. After transforming rice using the pMR285 multiplex CRISPR-Cas9 vector (Ritter et al., 2017) harboring those sgRNAs, we isolated T_0_ bi-allelic mutant lines containing various insertion and/or deletion mutations (indels) in the targeted *OsC4H1* and *OsC4H2a*/*2b* sites. The T_0_ bi-allelic mutants were further cultivated and screened to isolate homozygous mutant lines in the T_1_ and T_2_ generations. Accordingly, we successfully isolated T_1_ and T_2_ homozygous mutant lines for *OsC4H1*-single-knockout line (*OsC4H1*-KO) and *OsC4H2a*- and *OsC4H2b*-double-knockout line (*OsC4H2a*/*2b*-DKO), which harbor 1-bp insertions in the targeted *C4H* sites (**Figure 3A**). To generate *OsC4H1*-, *OsC4H2a*-, and *OsC4H2b*-triple-knockout lines, *OsC4H2a*/*2b*-DKO was further genome-edited by transforming the CRISPR-Cas9 vector targeting *OsC4H1*. Consequently, we successfully isolated two triple-knockout lines, *OsC4H1/2a*/*2b*-TKO-a and *OsC4H1/2a*/*2b*-TKO-b, harboring 1-bp insertions in the targeted *OsC4H1* site along with the *OsC4H2a*- and *OsC4H2b* sites (**Figure 3A**). All the indels resulted in frameshift mutations and the formation of premature stop codons, resulting in the loss of function of OsC4H1, OsC4H2a, and/or OsC4H2b (**Supplemental Figure S2–S4**). We also determined that there were no off-target mutations in the three top-ranked potential off-target sites in all the mutant plants used for further analysis (**Supplemental Table S4**).

**Figure 3.**
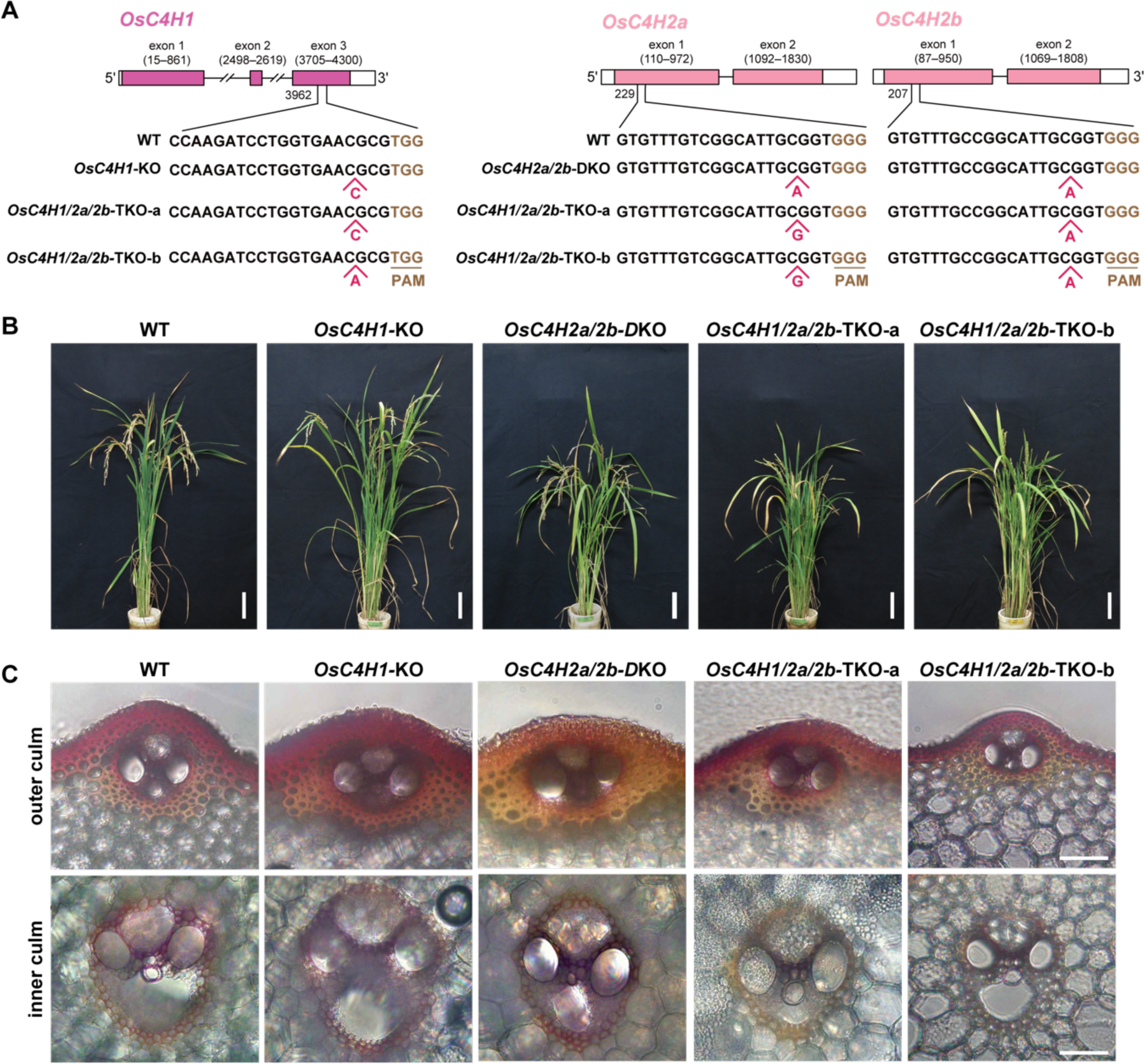
Genotype, phenotype and cell wall morphology of *C4H*-knockout rice. **A**) Gene structures and mutation patterns of *OsC4H1*, *OsC4H2a* and *OsC4H2b* loci in the *C4H*-knockout mutants generated by CRISPR/Cas9 editing. The protospacer adjacent motif (PAM) and inserted nucleotides in the sequences are highlighted. **B**) Images of wild-type and *C4H*-deficient plants at ripening stage. Bars = 10 cm. **C**) Histochemical analysis of outer (upper pictures) and inner (lower pictures) culm cell walls from wild-type and *C4H*-knockout plants at heading stage using the phloroglucinol-HCl lignin stain. Scale bars denote 50 µm. WT, wild type control line; *OsC4H1*-KO, *OsC4H1* single-knockout line; *OsC4H2a/2b*-DKO, *OsC4H2a* and *OsC4H2b* double-knockout line; *OsC4H1/2a/2b*-TKO-a and *OsC4H1/2a/2b*-TKO-b, *OsC4H1*, *OsC4H2a* and *OsC4H2b* triple-knockout lines.

### Phenotype and vascular anatomy of *C4H*-knockout rice mutants

The fully genotyped homozygotes of *OsC4H1*-KO, *OsC4H2a*/*2b*-DKO, *OsC4H1/2a*/*2b*-TKO-a and *OsC4H1/2a*/*2b*-TKO-b mutant lines (T_2_ generation) were grown side-by-side with the wild-type (WT) control line for phenotype and cell wall characterizations. At the 4-week-old seedling stage, all mutant lines displayed significantly reduced shoot height (by 8–22%), shoot biomass (by 19–24%), and root biomass (by 25–30%) (**Table 1**). At the 5-month-old mature stage (**Figure 3B**), all mutant lines still exhibited significantly reduced culm length (by 14–25%). Additionally, *OsC4H1/2a/2b*-TKO-a and *OsC4H1/2a/2b*-TKO-b showed reductions in plant height (by 9–12%) (**Table 1**). However, there were no significant differences in tiller number and aerial biomass between the mutant and WT control plants at the 5-month-old mature stage. Notably, *OsC4H1*-deficient mutant lines, i.e., *OsC4H1*-KO, *OsC4H1/2a/2b*-TKO-a, and *OsC4H1/2a/2b*-TKO-b, displayed considerable reductions in seed fertility (by 77–83%), whereas no significant reduction in seed fertility was detected in *OsC4H2a/2b*-DKO compared to WT (**Table 1**). We also examined the vascular anatomy and lignin deposition of the *C4H*-knockout rice mutants by analyzing their developing culms using the Wiesner (phloroglucinol-HCl) lignin staining solution. No obvious abnormalities in the overall vasculature (e.g., collapsed xylem cells and thinner secondary cell walls) were detected. However, slight reductions in the reddish coloration (i.e., lignin signal) from the cell walls were observed following staining with the Wiesner reagent (**Figure 3C**). Overall, these phenotype and cell wall morphology data of *C4H*-knockout rice mutants suggest that disruptions of *OsC4H1*, *OsC4H2a*, and *OsC4H2b* notably affect plant growth and lignin deposition. Nevertheless, it is remarkable that even the *OsC4H1/2a/2b*-TKO-a and *OsC4H1/2a/2b*-TKO-b mutants, which presumably lack any C4H activity, are still able to reach maturity and set seeds without significant reductions in biomass production. This sharply contrasts with previous C4H studies in eudicot model species (Schilmiller et al., 2009; El Houari et al., 2021; Kim et al., 2021; El Houari et al., 2021) as further discussed below.

**Table 1.**
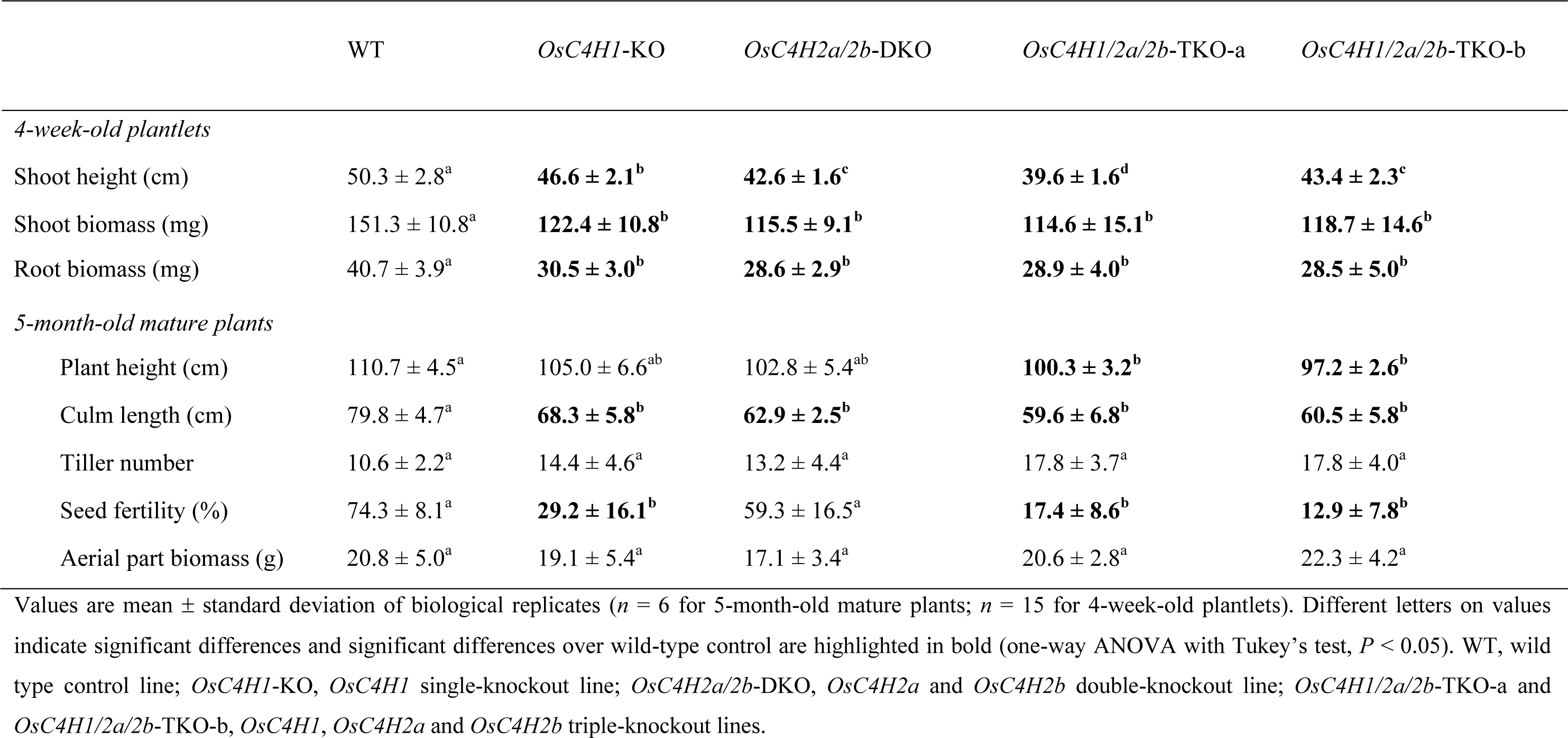
The growth performances of *C4H*-knockout rice mutants.

|  | WT | *OsC4H1*-KO | *OsC4H2a/2b*-DKO | *OsC4H1/2a/2b*-TKO-a | *OsC4H1/2a/2b*-TKO-b |
| --- | --- | --- | --- | --- | --- |
| *4-week-old plantlets* |  |  |  |  |  |
| Shoot height (cm) | 50.3 ± 2.8<sup>a</sup> | **46.6 ± 2.1<sup>b</sup>** | **42.6 ± 1.6<sup>c</sup>** | **39.6 ± 1.6<sup>d</sup>** | **43.4 ± 2.3<sup>c</sup>** |
| Shoot biomass (mg) | 151.3 ± 10.8<sup>a</sup> | **122.4 ± 10.8<sup>b</sup>** | **115.5 ± 9.1<sup>b</sup>** | **114.6 ± 15.1<sup>b</sup>** | **118.7 ± 14.6<sup>b</sup>** |
| Root biomass (mg) | 40.7 ± 3.9<sup>a</sup> | **30.5 ± 3.0<sup>b</sup>** | **28.6 ± 2.9<sup>b</sup>** | **28.9 ± 4.0<sup>b</sup>** | **28.5 ± 5.0<sup>b</sup>** |
| *5-month-old mature plants* |  |  |  |  |  |
| Plant height (cm) | 110.7 ± 4.5<sup>a</sup> | 105.0 ± 6.6<sup>ab</sup> | 102.8 ± 5.4<sup>ab</sup> | **100.3 ± 3.2<sup>b</sup>** | **97.2 ± 2.6<sup>b</sup>** |
| Culm length (cm) | 79.8 ± 4.7<sup>a</sup> | **68.3 ± 5.8<sup>b</sup>** | **62.9 ± 2.5<sup>b</sup>** | **59.6 ± 6.8<sup>b</sup>** | **60.5 ± 5.8<sup>b</sup>** |
| Tiller number | 10.6 ± 2.2<sup>a</sup> | 14.4 ± 4.6<sup>a</sup> | 13.2 ± 4.4<sup>a</sup> | 17.8 ± 3.7<sup>a</sup> | 17.8 ± 4.0<sup>a</sup> |
| Seed fertility (%) | 74.3 ± 8.1<sup>a</sup> | **29.2 ± 16.1<sup>b</sup>** | 59.3 ± 16.5<sup>a</sup> | **17.4 ± 8.6<sup>b</sup>** | **12.9 ± 7.8<sup>b</sup>** |
| Aerial part biomass (g) | 20.8 ± 5.0<sup>a</sup> | 19.1 ± 5.4<sup>a</sup> | 17.1 ± 3.4<sup>a</sup> | 20.6 ± 2.8<sup>a</sup> | 22.3 ± 4.2<sup>a</sup> |

Values are mean ± standard deviation of biological replicates (*n* = 6 for 5-month-old mature plants; *n* = 15 for 4-week-old plantlets). Different letters on values indicate significant differences and significant differences over wild-type control are highlighted in bold (one-way ANOVA with Tukey's test, *P* < 0.05). WT, wild type control line; *OsC4H1*-KO, *OsC4H1* single-knockout line; *OsC4H2a/2b*-DKO, *OsC4H2a* and *OsC4H2b* double-knockout line; *OsC4H1/2a/2b*-TKO-a and *OsC4H1/2a/2b*-TKO-b, *OsC4H1*, *OsC4H2a* and *OsC4H2b* triple-knockout lines.

### Cell wall structures of *C4H*-knockout rice mutants

To further investigate the impact of the loss-of-function mutations to the *C4H* genes on cell wall chemical structures, cell wall residue (CWR) samples prepared via a serial solvent extraction of the 5-month-old mature culm tissues of the *OsC4H1*-KO, *OsC4H2a/2b*-DKO, *OsC4H1/2a/2b*-TKO-a, and *OsC4H1/2a/2b*-TKO-b mutant lines (fully genotyped T_1_ homozygous individuals) as well as the WT control plants were subjected to in-depth cell wall structural analyses using chemical and nuclear magnetic resonance (NMR) methods. For lignin content analysis using the thioglycolic lignin assay, we also prepared CWR samples from the roots of hydroponically grown 4-week-old plantlets.

### Lignin content

The lignin content of the *C4H*-knockout rice mutant cell walls was evaluated on the basis of lignin levels determined by thioglycolic lignin assay (**Figure 4A**) as well as the changes in the total yields of the lignin-derived monomeric compounds released by thioacidolysis (**Figure 4C**) and derivatization followed by reductive cleavage (DFRC) (**Figure 4D**). In the culm, the *OsC4H1*-KO, *OsC4H1/2a/2b*-TKO-a, and *OsC4H1/2a/2b*-TKO-b mutants exhibited reduced lignin levels by 23–32% based on thioglycolic acid lignin assay (albeit not statistically significant for *OsC4H1/2a/2b*-TKO-a) (**Figure 4A**), by 24–30% based on thioacidolysis-derived product yield (**Figure 4C**), and by 28–37% based on DFRC-derived product yield (**Figure 4D**), compared to the WT control. In contrast, no statistically significant difference in lignin levels was detected between *OsC4H2a/2b*-DKO and the WT control. On the other hand, in the root, thioglycolic acid lignin assay detected lignin reductions in *OsC4H1/2a/2b*-TKO-a (by ∼20%) and *OsC4H1/2a/2b*-TKO-b (by ∼18%) but no significant change in *OsC4H1*-KO and *OsC4H2a/2b*-DKO (**Figure 4A**). These data suggest that, in alignment with the gene expression profiles of the *C4H* genes (**Figure 2**), OsC4H1 contributes to lignin biosynthesis in the culm, and OsC4H1, OsC4H2a, and OsC4H2b redundantly contribute to lignin biosynthesis in the root. Nevertheless, it should be emphasized again that the *OsC4H1/2a/2b*-TKO-a and *OsC4H1/2a/2b*-TKO-b mutants are still able to produce a substantial amount of lignin in both culm and root.

**Figure 4.**
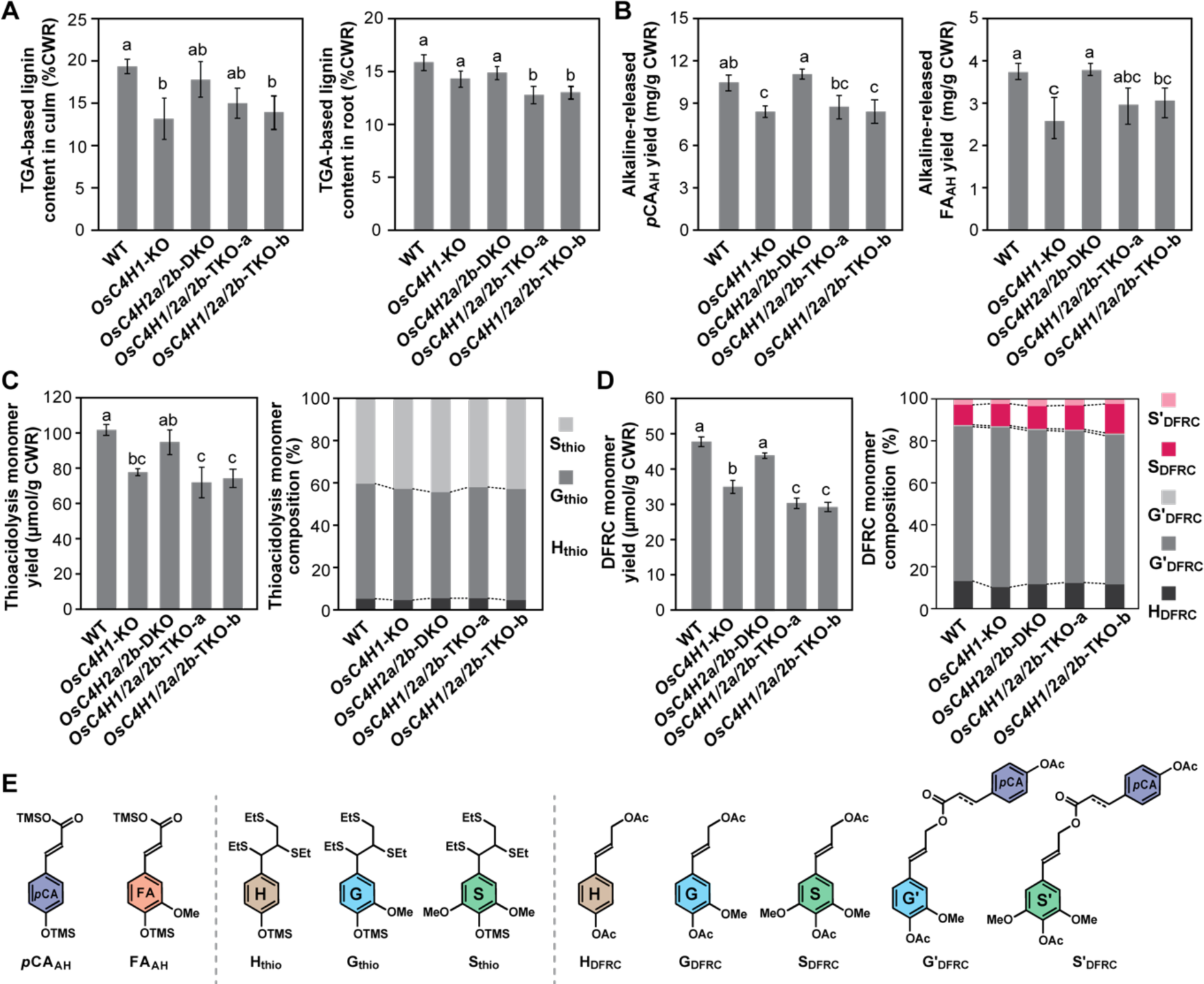
Lignin and hydroxycinnamate analysis of *C4H*-knockout rice cell walls. **A**) Lignin content in culm and root as determined by thioglycolic acid (TGA) lignin assay. **B**) Yield of *p*-coumarate (*p*CA) and ferulate (FA) released by alkaline hydrolysis. **C** and **D**) Yield and ratio of lignin degradation monomers released via thioacidolysis (**C**) and derivatization followed by reductive cleavage, DFRC (**D**). **E**) Structures of ***p*CA_AH_** and **FA_AH_** released by alkaline hydrolysis, **H_thio_**, **G_thio_** and **S_thio_** released by thioacidolysis (and silylated), and **H_DFRC-OH_**, **G_DFRC-OH_**, **S_DFRC-OH_**, **G_DFRC-pCA_**, and **S_DFRC-pCA_** released by DFRC (and acetylated). Analyses were performed on culm cell wall residue (CWR) samples prepared from dried mature rice plants or root CWR prepared from 4-week-old seedlings. Values are mean ± standard deviation of biological replicates (*n* = 3). Different letters on values indicate significant differences and significant differences over wild-type control (one-way ANOVA with Tukey’s test, *P* < 0.05). WT, wild type control line; *OsC4H1*-KO, *OsC4H1* single-knockout line; *OsC4H2a/2b*-DKO, *OsC4H2a* and *OsC4H2b* double-knockout line; *OsC4H1/2a/2b*-TKO-a and *OsC4H1/2a/2b*-TKO-b, *OsC4H1*, *OsC4H2a* and *OsC4H2b* triple-knockout lines.

### Lignin composition

Thioacidolysis and DFRC were also used to determine the lignin compositional changes in the *C4H*-knockout rice mutant cell walls. Thioacidolysis releases thioethylated H- (**H_thio_**), G- (**G_thio_**), and S-type (**S_thio_**) monomeric compounds from the H-, G-, and S-type β–*O*–4 units, respectively, regardless of their γ-acylation status (Lapierre et al., 1986) (**Figure 4E**). On the other hand, DFRC releases quantifiable amounts of non-γ-acylated (γ-free) monomeric compounds (**H_DFRC_**, **G_DFRC_**, and **S_DFRC_**) and γ-*p*-coumaroylated monomeric compounds (**Gʹ_DFRC_**and **Sʹ_DFRC_**) from the corresponding non-γ-acylated and γ-*p*-coumaroylated β–*O*–4 units (Lu and Ralph, 1997, 1999; Karlen et al., 2018) (**Figure 4E**). Consequently, neither thioacidolysis nor DFRC detected any clear difference in lignin composition between all the examined *C4H*-knockout mutants and WT control cell walls (**Figure 4C and 4D**).

### Cell wall-bound hydroxycinnamates and polysaccharides

The levels of cell wall-bound hydroxycinnamates, i.e., *p*-coumarate (*p*CA) mainly attached on lignin and ferulate (FA) mainly attached on arabinoxylan (Ralph, 2010), in the rice culm cell walls, were evaluated by quantifying the *p*CA (***p*CA_AH_**) and FA (**FA_AH_**) (**Figure 4E**) released via a mild alkaline treatment of the CWR samples. Similar to the changes in lignin levels, both ***p*CA_AH_** and **FA_AH_**levels were reduced in the *OsC4H1*-KO (by ∼20% and ∼31% for ***p*CA_AH_**and **FA_AH_**, respectively), *OsC4H1/2a/2b*-TKO-a (by ∼17% and ∼16% for ***p*CA_AH_** and **FA_AH_**, respectively, albeit no statistical significance), and *OsC4H1/2a/2b*-TKO-b (by ∼20% and ∼18% for ***p*CA_AH_** and **FA_AH_**, respectively) mutants, but not in *OsC4H2a/2b*-DKO (**Figure 4B**). These data suggested that, similar to its proposed role in the biosynthesis of monolignol-derived lignin units, OsC4H1 contributes to the biosynthesis of cell wall-bound *p*CA and FA in rice culm.

We also investigated the abundance of cell wall polysaccharides by performing a neutral sugar analysis according to the two-step acid-catalyzed hydrolysis method. Overall, the abundance of neutral sugars released from the *C4H*-knockout rice culm cell walls was similar to those from the WT control, although notable increases in the amounts of amorphous glucose (glucose released from trifluoracetic acid-soluble fractions) were detected in *OsC4H1*-KO, *OsC4H2a/2b*-DKO, and *OsC4H1/2a/2b*-TKO-a (**Supplemental Table S5**).

### 2D HSQC NMR

To further characterize the cell wall structures of the *C4H*-knockout rice mutants, we conducted a solution-state two-dimensional (2D) ^1^H–^13^C heteronuclear single-quantum coherence (HSQC) NMR analysis on the culm cell walls prepared from the 5-month-old mature *OsC4H1*-KO, *OsC4H2a/2b*-DKO, *OsC4H1/2a/2b*-TKO-a, and *OsC4H1/2a/2b*-TKO-b mutant and WT control plants. The 2D HSQC NMR spectra of the whole cell walls were collected by directly swelling ball-milled rice CWR samples in the dimethyl sulfoxide (DMSO)-*d*_6_/pyridine-*d*_5_ NMR solvent system (Kim and Ralph, 2010; Mansfield et al., 2012). The obtained rice cell wall spectra displayed contour signals from typical monolignol-derived lignin (**S**, **G**, and **H**), lignin-bound tricin (**T**), and cell wall-bound hydroxycinnamates (***p*CA** and **FA**), along with signals from cell wall polysaccharides typical of grass cell walls (**Gl**, **Xy**, **Xyʹ**, **Xyʹʹ**, **Ar**, **Ga**, and **GlU**) (**Supplemental Figure S5** and **Supplemental Table S6**). For a semi-quantitative examination of the relative proportions of lignin and polysaccharide unis in the rice cell walls, we performed volume integration analysis of the well-resolved, major lignin, *p*-hydroxycinnamate, and polysaccharide signals (**Figure 5**). The reported signal intensity data are relative intensities normalized according to the sum of the integrated signals, reflecting the proportional amount of each component in the rice cell walls (see **Materials and Methods**).

**Figure 5.**
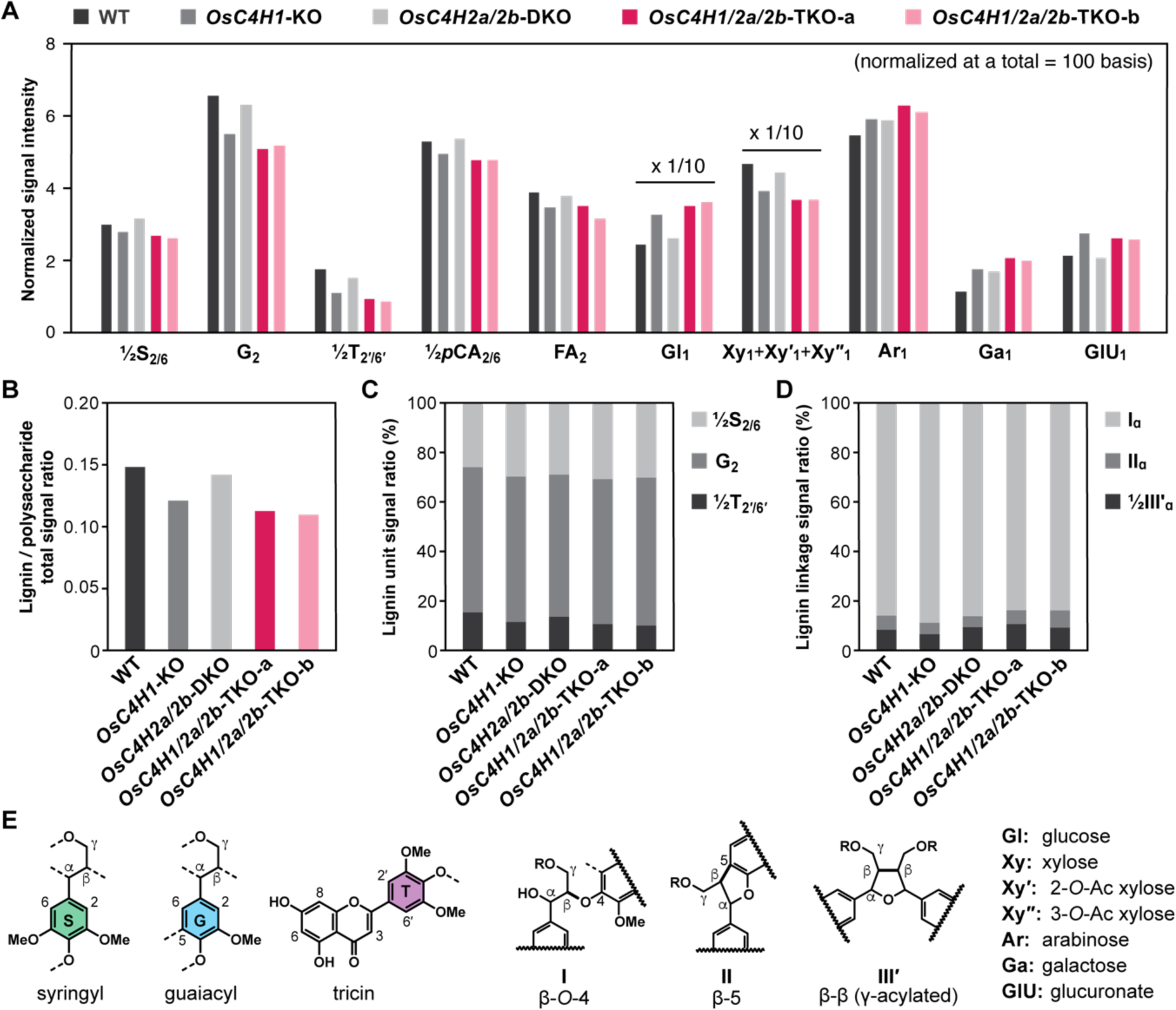
2D HSQC NMR signal intensity analysis of *C4H*-knockout rice culm cell walls. **A**) Relative signal intensity of major lignin, hydroxycinnamate and polysaccharide signals normalized based on the sum of the integrated signals (**½S_2/6_** + **G_2_** + **½T_2ʹ/6ʹ_** + **½P_2/6_** + **FA_2_** + **Gl_1_** + **Xy_1_** + **Xyʹ_1_** + **Xyʹʹ_1_** + **Ar_1_** + **Ga_1_** + **GlU_1_**). Data labeled ×1/10 denote data that has been divided by a factor 10 for visualization. **B**) Signal intensity ratio of major lignin (**½S_2/6_** + **G_2_** + **½T_2ʹ/6ʹ_**) and polysaccharide (**Gl_1_** + **Xy_1_** + **Xyʹ_1_** + **Xyʹʹ_1_** + **Ar_1_**) signals. **C**) Signal intensity ratio for syringyl (**½S_2/6_**), guaiacyl (**G_2_**), and tricin (**½ T_2ʹ/6ʹ_**) lignin units. **D**) Signal intensity ratio for major lignin inter-monomeric linkages (**I_ɑ_**, **II_ɑ_**, and **½III′_ɑ_**). **E**) Structures of major lignin aromatic units and inter-monomeric linkages. NMR spectra and complete signal assignments are shown in **Supplemental Figure S5** and **Supplemental Table S6**. NMR analysis was conducted for culm cell wall residue (CWR) samples pooled from three biologically independent plants for each line. WT, wild type control line; *OsC4H1*-KO, *OsC4H1* single-knockout line; *OsC4H2a/2b*-DKO, *OsC4H2a* and *OsC4H2b* double-knockout line; *OsC4H1/2a/2b*-TKO-a and *OsC4H1/2a/2b*-TKO-b, *OsC4H1*, *OsC4H2a* and *OsC4H2b* triple-knockout lines.

In the cell wall spectra of the *OsC4H1*-KO, *OsC4H1/2a/2b*-TKO-a, and *OsC4H1/2a/2b*-TKO-b mutants, relative signal intensities of the monolignol-derived G- and S-type lignin aromatic units (**G** and **S**) as well as cell-wall-bound hydroxycinnamates (***p*CA** and **FA**) were reduced compared to those observed in the wild-type control spectrum (**Figure 5A**), further supporting that OsC4H1 contributes to the biosynthesis of monolignol-derived lignin units as well as cell-wall-bound *p*CA and FA in rice culm; the signals from H-type aromatic units (**H**) were excluded from the volume integration analysis because of their relatively low abundance (**Figure 4**) and their signal overlap with protein-derived aromatic signals (Kim et al., 2017). In addition, we detected similar reductions in the signal intensities of lignin-bound tricin (**T**) in the cell wall spectra of the *OsC4H1*-KO, *OsC4H1/2a/2b*-TKO-a, and *OsC4H1/2a/2b*-TKO-b mutants (**Figure 5A**), indicating that OsC4H1 contributes to the biosynthesis of lignin-bound tricin in rice culm. These aromatic signals were also slightly reduced in the cell wall spectra of the *OsC4H2a/2b*-DKO mutant but less prominently compared to the reductions observed in the spectra of *OsC4H1*-knockout mutants (*OsC4H1*-KO, *OsC4H1/2a/2b*-TKO-a, and *OsC4H1/2a/2b*-TKO-b) (**Figure 5A**). Consequently, the signal intensity ratios of the major lignin aromatic (**½S_2/6_** + **G_2_** + **½T_2ʹ/6ʹ_**) to polysaccharide anomeric (**Gl_1_** + **Xy_1_** + **Xyʹ_1_**+ **Xyʹʹ_1_** + **Ar_1_**) signals were notably reduced in the spectra of the *OsC4H1*-KO, *OsC4H1/2a/2b*-TKO-a, and *OsC4H1/2a/2b*-TKO-b mutants, but less prominently in the spectrum of the *OsC4H2a/2b*-DKO mutant (**Figure 5B**). No clear change in the proportions of the G, S, and tricin lignin signals (**G**, **S**, and **T**) was found in the cell wall spectra of all examined *C4H*-knockout rice mutants (**Figure 5C**). In addition, we detected no clear change in the distribution of the major lignin inter-monomeric linkages, i.e., β–*O*–4 (**I**), β–5 (**II**), and β–β (**IIIʹ**), in the cell wall spectra of all examined *C4H*-knockout rice mutants (**Figure 5D**).

Overall, the chemical and NMR structural data of the *C4H*-knockout rice mutants demonstrate the roles of OsC4H1, OsC4H2a, and OsC4H2b in the biosynthesis of lignin building blocks, such as monolignols, tricin, and hydroxycinnamates (*p*CA and FA), contributing to cell wall development in rice. However, concurrently, these data clearly indicate that rice can biosynthesize these lignin building blocks and deposit substantial levels of lignin to develop reasonably robust cell walls even without C4H function, as further investigated below.

### Contributions of the dual PAL-C4H and PTAL pathways

The less prominent impacts of the loss of C4H function in rice may be attributed to the presence of the parallel PTAL pathway, which can supply lignin precursors from Tyr in addition to the conventional PAL-C4H pathway from Phe (**Figure 1**) (Barros et al., 2016; Barros and Dixon, 2020). To address this, we further characterized the *C4H*-knockout rice mutants along with wild-type rice using ^13^C-isotope labeling and chemical inhibitor assays.

### Feeding experiment using ^13^C-labeled Phe and Tyr

To assess the capability of rice to incorporate Phe and Tyr into lignin, we performed feeding experiments using ^13^C-labeled Phe and Tyr on the WT and *OsC4H1/2a/2b*-TKO-a mutant seedlings. Rice seedlings were grown in a half Murashige and Skoog (MS) medium supplemented with 1 mM ^13^C_6_-labeled Phe or ^13^C_9_-labeled Tyr for 30 days. CWR samples prepared from the shoot parts of the fed seedlings were then subjected to thioacidolysis and alkaline hydrolysis (**Supplemental Figure S6 and S7**). The resulting monomeric products released from monolignol-derived lignin units (**H_thio_**, **G_thio_**, and **S_thio_**), tricin-lignin units (**T_Thio_**) and cell wall-bound *p*CA and FA (***p*CA_AH_** and **FA_AH_**) were analyzed by gas chromatography-mass spectrometry (GC-MS) (for **H_thio_**, **G_thio_**, **S_thio_**, ***p*CA_AH_**, and **FA_AH_**) or liquid chromatography-mass spectrometry (LC-MS) (for **T_thio_**) to assess the incorporation of ^13^C-labeled Phe and Tyr into each of the lignin and hydroxycinnamate units (**Supplemental Figure S6 and S7**).

As shown in **Figure 6**, the capability of the WT rice seedlings to incorporate ^13^C-labeled Phe and Tyr into lignin and hydroxycinnamate units was clearly demonstrated. We observed higher incorporations of ^13^C-labeled Phe than ^13^C-labeled Tyr into all the examined lignin and hydroxycinnamate units, with the exception of cell wall-bound FA (**FA_AH_**), where comparable levels of ^13^C-labeled Phe and Tyr were detected. In contrast, the *OsC4H1/2a/2b*-TKO-a mutant seedlings showed no detectable incorporation of ¹³C-labeled Phe, while the incorporation levels of ¹³C-labeled Tyr into all the examined lignin and hydroxycinnamate units remained comparable to or slightly exceeded those observed in the WT seedlings (**Figure 6**). This result affirms that the *OsC4H1/2a/2b*-TKO-a mutant completely lacks C4H activity and demonstrates that rice can still produce all the major lignin and hydroxycinnamate precursors, i.e., H-, G-, and S-type monolignols, tricin, *p*CA, and FA, for cell wall lignification from Tyr even without C4H activity.

**Figure 6.**
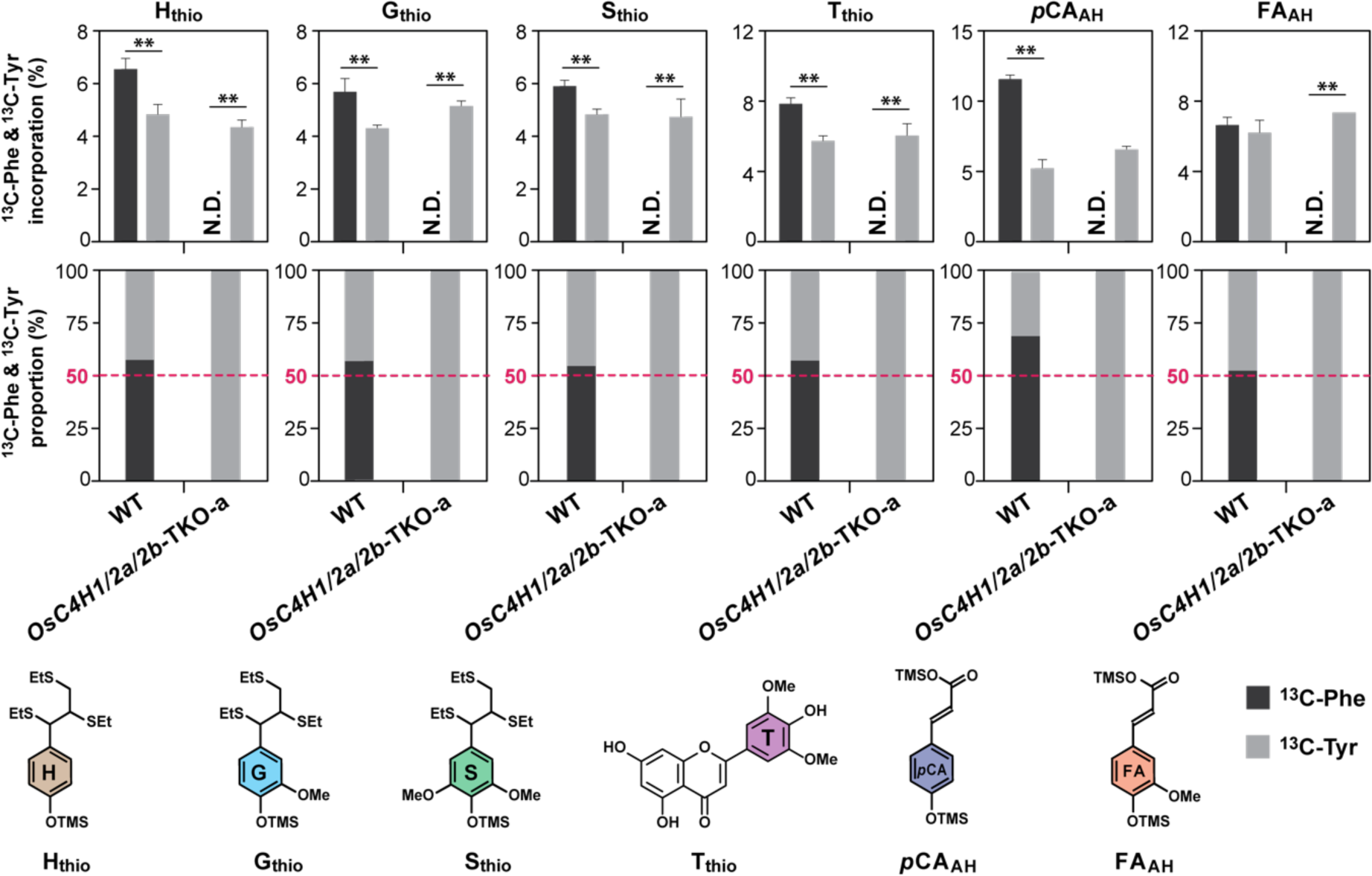
Incorporation of ^13^C-labeled phenylalanine and tyrosine into lignin polymer units in wild-type and *C4H*-knockout mutant rice seedlings. Incorporation rates (upper) and proportions (lower) of ^13^C-labeled phenylalanine (Phe) and tyrosine (Tyr) into lignin and hydroxycinnamate units released from shoot cell walls of wild-type and *C4H*-knockout mutant rice. **C**) Structures of thioacidolysis-derived lignin monomeric products (**H_thio_**, **G_thio_**, **S_thio_** and **T_thio_**) and alkaline hydrolysis-derived hydroxycinnamates (***p*CA_AH_** and **FA_AH_**). Rice seedlings were grown in a medium containing 1 mM ^13^C_6_-labeled Phe and ^13^C_9_-labeled Tyr for 30 days. Cell wall samples prepared from the shoot parts of the fed seedlings were subjected to thioacidolysis and alkaline hydrolysis. The released products were analyzed by GC- or LC-MS to determine the ^13^C incorporation rate. All analyses were performed in triplicate on pooled cell wall residue (CWR) samples collected from 25-30 plants. Asterisks indicate statistically significant differences (Student’s *t*-test, *n* = 3, \*\**p* < 0.01). N.D., not detected. WT, wild-type control line; *OsC4H1/2a/2b*-TKO-a, *OsC4H1*, *OsC4H2a* and *OsC4H2b* triple-knockout line.

### Chemical inhibitor assays using C4H and PAL/TAL inhibitors

To further examine the contributions of the PAL-C4H and PTAL pathways in wild-type and *C4H*-knockout rice, we employed chemical inhibitor assays using piperonylic acid (PA) (Schalk et al., 1998) and L-2-aminooxy-3-phenylpropionic acid (AOPP) (Holländer et al., 1979) as C4H and PAL/TAL inhibitors, respectively; while AOPP has long been used as a PAL inhibitor, earlier reports showed that it also exhibits a strong inhibitory effect on TAL activities when applied to grasses (Holländer et al., 1979; Rösler et al., 1997). The 5-day-old seedlings of the WT, *OsC4H1/2a/2b*-TKO-a and *OsC4H1/2a/2b*-TKO-b lines were treated in a hydroponic solution containing PA (1 mM) and/or AOPP (3 mM) for 9 days, and then subjected to phenotypic characterization and lignin analysis of shoot CWR using thioacidolysis.

The WT seedlings treated with 1 mM PA exhibited significant reductions in shoot length under the tested conditions, albeit with no significant reduction in shoot biomass and primary root length and root biomass compared to the mock control (**Figure 7A**). The PA-treated WT seedlings also exhibited a decrease in the amounts of thioacidolysis-derived lignin degradation products in shoot CWRs (**Figure 7B**). These impaired plant growth and lignin deposition in the WT seedlings can be primarily attributed to the inhibition of the PAL-C4H pathway by PA. In line with this notion, the PA treatment did not affect the growth and lignin deposition in the *OsC4H1/2a/2b*-TKO-a and *OsC4H1/2a/2b*-TKO-b seedlings in which the PAL-C4H pathway is completely blocked (**Figure 7A and 7B**). The application of 3 mM AOPP to block both PAL-C4H and PTAL pathways more prominently reduced most of the growth parameters examined (all except primary root length) than the PA treatment in the WT seedlings (**Figure 7A**), whereas the amounts of thioacidolysis-derived lignin degradation products from the AOPP-treated WT seedlings decreased compared to the mock control but remained similar to the PA-treated WT seedlings (**Figure 7B**). The AOPP treatment also significantly reduced the growth parameters and the amounts of thioacidolysis-derived lignin degradation products in the *OsC4H1/2a/2b*-TKO-a and *OsC4H1/2a/2b*-TKO-b seedlings (**Figure 7A and 7B**). Adding 1 mM PA along with 3 mM AOPP did not cause any further substantial reduction in either growth parameters or the amounts of lignin degradation products in all the tested rice lines (**Figure 7A and 7B**). Overall, these results further support the notion that the parallel PAL-C4H and PTAL pathways cooperatively contribute to lignification in rice.

**Figure 7.**
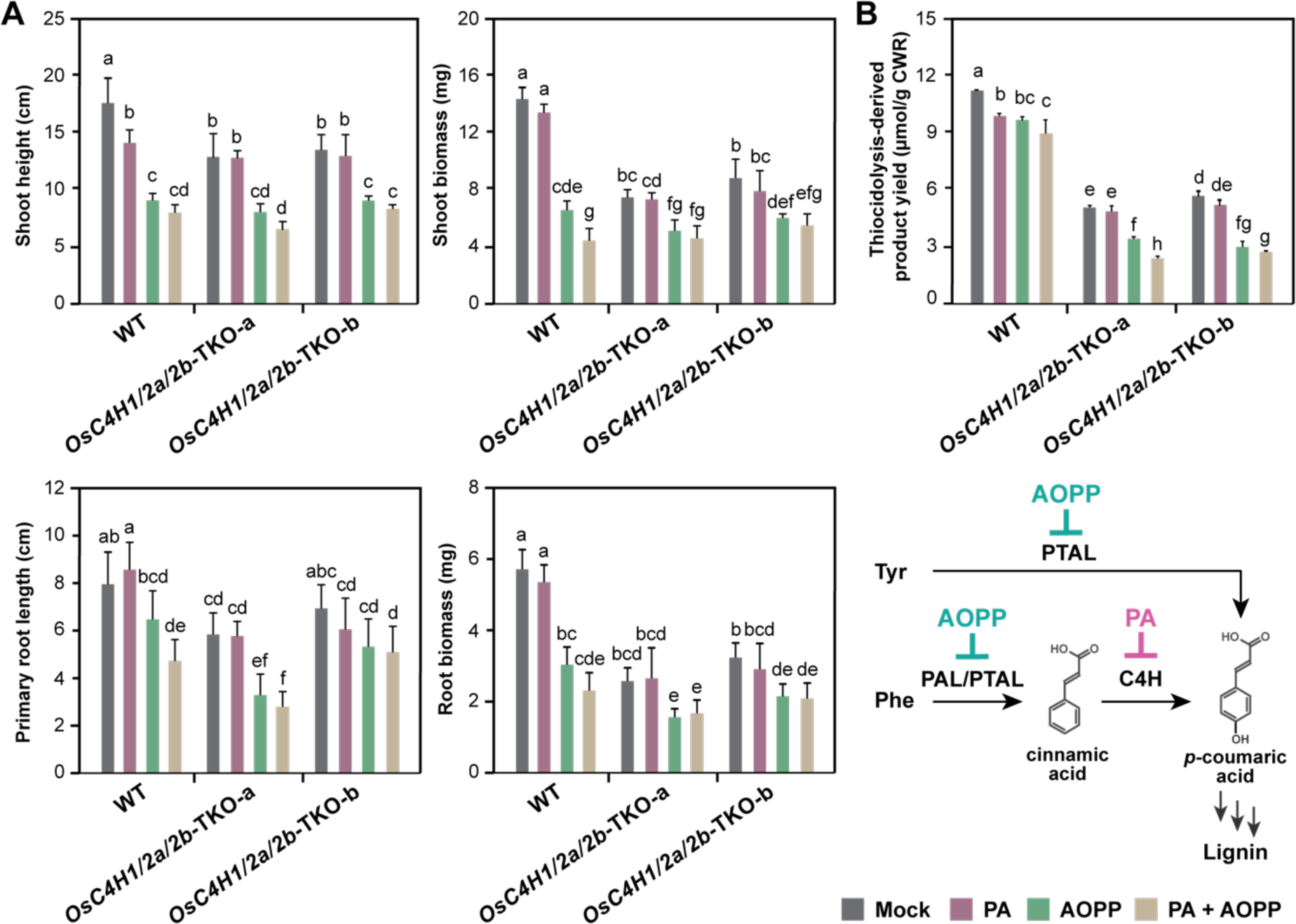
Chemical inhibitor assay of wild-type and *C4H*-knockout mutant rice seedlings. **A**) Growth parameters of the WT and *C4H*-knockout mutant rice seedlings treated with piperonylic acid (PA) and/or L-2-aminooxy-3-phenylpropionic acid (AOPP). Values are mean ± standard deviation of biological replicates (*n* = 8). **B**) Total yield of thioacidolysis-derived monomeric products (**H_thio_** + **G_thio_** + **S_thio_**) released from shoot cell wall residue (CWR) samples. Analysis was conducted for three sets of pooled shoots CWR samples prepared from 2-3 plants for each (*n* = 3). Different letters indicate significant differences by one-way ANOVA with Tukey’s test (*P* < 0.05). WT, wild type control line; *OsC4H1/2a/2b*-TKO-a and *OsC4H1/2a/2b*-TKO-b, *OsC4H1*, *OsC4H2a* and *OsC4H2b* triple-knockout lines.

## Discussion

Extensive research has elucidated C4H’s function in eudicot lignin biosynthesis, but its role in grasses remains poorly understood due to a scarcity of knockout mutants. In particular, the role of C4H in the presence of the grass-specific Tyr-derived PTAL pathway alongside the conserved Phe-derived PAL-C4H pathway remains elusive. This study significantly addresses this gap by characterizing a set of rice mutants with single and multiple knockouts in major *C4H* genes generated through CRISPR-Cas9 genome editing.

### Class I and class II C4H isoforms cooperatively contribute to lignin biosynthesis in rice

Most vascular plants, including gymnosperms and angiosperms (except eudicot Brassicaceae such as Arabidopsis), possess at least two copies of C4H paralogs, namely class I and class II C4H isoforms (Renault et al., 2017; Renault et al., 2019; Knosp et al., 2024). While the role of class I C4Hs in lignin biosynthesis has been well-documented in eudicots, the involvement of class II C4Hs in lignin biosynthesis remains obscure even in eudicots (Renault et al., 2017). An earlier study reported that antisense suppression of class II C4H in transgenic tobacco resulted in notable reductions in cell wall lignin (Blee et al., 2001). In addition, both class I (CYP73A92 and CYP73A93) and class II (CYP73A94) C4H proteins from *B. distachyon* **(Supplemental Table S1**) have been shown to exhibit typical C4H enzymatic activity *in vitro* and be capable of complementing *C4H*-deficient Arabidopsis mutants (Renault et al., 2017; Knosp et al., 2024).

The functional characterization of rice mutants with targeted knockouts in *C4H* genes encoding class I [OsC4H1 (CYP73A38)] and class II [OsC4H2a (CYP73A39) and OsC4H2b (CYP73A40)] proteins strongly supports their cooperative roles in lignin biosynthesis. Notably, *OsC4H1* expression levels were significantly higher than those of *OsC4H2a* and *OsC4H2b* in major vegetative organs of the aerial parts (culm, leaf blade, and leaf sheath) (**Figure 2**). Consistent with this expression pattern, both *OsC4H1* knockout (*OsC4H1*-KO) and triple knockout (*OsC4H1/2a/2b*-TKO) plants displayed similar reductions in lignin content within culm cell walls, whereas, in contrast, the double knockout of *OsC4H2a* and *OsC4H2b* (*OsC4H2a/2b*-DKO) showed no significant change in lignin levels compared to wild-type controls (**Figure 4 and 5**). Our chemical and 2D NMR analyses revealed reductions not only in monolignol-derived lignin units but also in tricin and hydroxycinnamate (*p*CA and FA) units associated with lignin and/or hemicelluloses in the culm cell walls of *OsC4H1*-KO and *OsC4H1/2a/2b*-TKO plants. These data collectively demonstrate that OsC4H1, but not OsC4H2a and OsC4H2b, plays the predominant role as C4H in culm lignification.

In contrast to the culm, *OsC4H1*, *OsC4H2a*, and *OsC4H2b* displayed comparable expression levels in the root (**Figure 2**). In line with these gene expression profiles, only the triple knockout (*OsC4H1/2a/2b*-TKO-a and *OsC4H1/2a/2b*-TKO-b) plants exhibited significant reductions in root cell wall lignin, while the independent knockouts of class I (*OsC4H1*-KO) and class II (*OsC4H2a/2b*-DKO) *C4H* genes did not **(Figure 4**). These data suggest that the two class II C4Hs (OsC4H2a and OsC4H2b) likely act cooperatively with the class I C4H (OsC4H1) to contribute to root lignification. Despite the apparent redundancy in rice roots, the high degree of conservation of both class I and class II C4H paralogs across vascular plants implies distinct and essential functions for each paralog (Renault et al., 2017; Renault et al., 2019). To further elucidate these potential functional specializations in rice, more rigorous analyses are needed, including further biochemical characterizations of class I and class II C4H proteins, and their gene expression and metabolite profiling at tissue and cellular levels.

### Distinct impacts of *C4H*-deficiency on plant development and lignin biosynthesis in rice versus eudicot species

The complete absence of C4H activity in *OsC4H1/2a/2b*-TKO plants was confirmed by their inability to incorporate ^13^C-labeled Phe into cell wall lignin and hydroxycinnamate units (**Figure 6**). This functional disruption indeed resulted in significant growth defects and reduced lignin levels in the cell walls of *OsC4H1/2a/2b*-TKO rice compared to wild-type controls (**Figure 3-5 and Table 1**). Nevertheless, the impact on plant growth and lignin biosynthesis in these *C4H*-nulll rice mutants appeared considerably less severe than what has been reported in *C4H*-deficient eudicots. *OsC4H1/2a/2b*-TKO rice developed to maturity with no significant reductions in shoot biomass. Albeit with largely decreased fertility, they set seeds (**Table 1**). No apparent cell wall morphological defects were observed using histochemical analysis of *OsC4H1/2a/2b*-TKO rice (**Figure 3**). Furthermore, the lignin reduction in these mutant plants was modest, reaching a maximum of 30% reduction in culms and 20% reduction in roots (**Figure 4-6**). These relatively mild growth and lignin biosynthesis defects in the *C4H*-null rice mutants stand in stark contrast to observations in *C4H*-deficient Arabidopsis *ref3* mutants. Complete C4H inactivation in *ref3* mutants results in seed lethality, while even leaky residual *C4H* expression leads to severe growth defects (abnormal stem and leaf development, collapsed vasculature, male sterility, and dwarfism) and substantial lignin reductions, reaching up to 80% in surviving plants (Schilmiller et al., 2009; El Houari et al., 2021; Kim et al., 2021). Overall, this study revealed that C4H activity is largely dispensable for plant growth and lignin biosynthesis in rice. This observation likely arises from the presence of the grass-specific PTAL pathway in rice, which offers an alternative route for lignin monomer and other phenylpropanoid synthesis, bypassing the need for the canonical PAL-C4H pathway, as further discussed below.

### The Tyr-derived PTAL pathway can largely compensate for the disruption of the Phe-derived PAL-C4H pathway in rice

Earlier studies using radioisotope labeling approaches provided the first evidence for TAL activity directing Tyr into phenylpropanoid biosynthesis in grasses (Brown and Neish, 1956; Neish, 1961; Jangaard, 1974). This finding was later solidified through biochemical and genetic characterization of grass-specific PTAL enzymes from maize (*Zea mays*) (Rösler et al., 1997), Brachypodium (Cass et al., 2015; Barros et al., 2016), and sorghum (*Sorghum bicolor*) (Jun et al., 2018), as comprehensively reviewed by Barros and Dixon (2020). A recent study using ^13^CO_2_ feeding estimated that Brachypodium, a model grass, can produce Tyr at rates exceeding ten times that of the eudicot model Arabidopsis, without impacting Phe biosynthesis (El-Azaz et al., 2023). Furthermore, MS-based analyses of lignin from Brachypodium seedlings fed with ^13^C-labeled Phe and Tyr revealed that a significant portion of lignin units in the cell walls originate from Tyr via the PTAL pathway, with additional contributions from Phe through the canonical PAL-C4H pathway (Barros et al., 2016; Barros et al., 2022). Likewise, this study demonstrates that both wild-type and *OsC4H1/2a/2b*-TKO rice plants incorporate ^13^C-labeled Tyr into all the major lignin and hydroxycinnamate units, including H-, G-, and S-type monolignol-derived lignin units, tricin-lignin units, and cell wall-bound *p*CA and FA units, in the cell walls (**Figure 6**). Based on the relative incorporation rates of ^13^C-labeled Phe and Tyr supplied to wild-type rice seedlings, we estimate that approximately 40–50% of the major lignin and hydroxycinnamate units in rice cell walls originate from Tyr via the PTAL pathway with the remaining 50–60% from Phe via the PAL-C4H pathway. As expected, *OsC4H1/2a/2b*-TKO rice plants, lacking a functional PAL-C4H pathway, rely solely on the PTAL pathway for Tyr incorporation into their major lignin and hydroxycinnamate units (**Figure 6**). Furthermore, chemical inhibition assays revealed that, at least under the tested conditions, rice displayed a significantly lower sensitivity to the C4H inhibitor (PA) compared to the PAL/PTAL inhibitor (AOPP) (**Figure 7**). Collectively, these findings support the view that the grass-specific PTAL pathway, bypassing the canonical C4H-PAL pathway, is operational in rice and can largely compensate for the reduction in lignin and associated phenylpropanoid biosynthesis when the C4H-PAL pathway is completely blocked.

Although the PTAL pathway significantly compensates for the disrupted PAL-C4H pathway in *C4H*-deficient rice mutants, the observed reductions in growth and lignin deposition in our *C4H*-deficient rice mutants suggest that metabolite pools from the two pathways are not entirely shared or equivalent. Despite observing subtle changes in lignin and hydroxycinnamate composition in our *C4H*-deficient rice mutants (**Figure 4 and 5**), and overall similar Phe/Tyr incorporation capacities among different lignin and hydroxycinnamate units in the wild-type control **(Figure 6)**, the Phe-derived and Tyr-derived *p*-coumaric acid generated by each pathway maybe, at least partially, directed towards different downstream products and/or targeted to different locations at different developmental stages. In line with this notion, prior isotope labeling studies of grass lignin and associated metabolites revealed variations in the incorporation ratios of Phe and Tyr within different types of lignin subunits (Barros et al., 2016; Barros et al., 2022) and soluble phenylpropanoids (Simpson et al., 2021). These findings warrant further in-depth studies on metabolites, genes, and enzymes associated with both pathways, at tissue and cellular levels. Unraveling the underlying mechanisms may provide valuable genetic tools for engineering grass cell walls and aromatic metabolisms. This, in turn, can facilitate the production and utilization of grass biomass and biochemicals for various applications.

## Materials and Methods

### Bioinformatics

The unrooted phylogenetic tree was constructed using the maximum-likelihood method in MEGA X (Kumar et al., 2018). Bootstrapping with 1,000 replications was performed. Microarray-based gene expression data were retrieved from the Rice Expression Profile Database (Sato et al., 2010). Gene locus and accession numbers of C4H genes and proteins examined are listed in **Supplemental Table S1**.

### Gene expression analysis by RT-qPCR

Total RNA was extracted from the culm, leaf blade, and leaf sheath at the heading stage and from the root at the 2-month-old seedling stage of rice as described previously (Koshiba et al. 2013). RNA was reverse-transcribed into cDNA using random hexamer primers (Invitrogen, Carlsbad, CA, USA) (Koshiba et al. 2013). Gene expression was assayed using an Applied Biosystems Step One Real-time PCR System (Applied Biosystems, Forster City, CA, USA) and primer sets listed in **Supplemental Table S7**. A ubiquitin gene (*OsUBQ5*; AK061988) was used as an internal control.

### Generation of rice mutants by CRISPR-Cas9

The *OsC4H1*-KO, *OsC4H2a/2b*-DKO, *OsC4H1/2a/2b*-TKO-a, and *OsC4H1/2a/2b*-TKO-b mutant lines were generated via CRISPR-Cas9-mediated mutagenesis using the pMR285 binary vector (Ritter et al., 2017) as described in **Supplemental Methods**. Selected T_1_ and T_2_ mutant lines used for the phenotypic characterization and cell wall analyses were grown to maturity in a greenhouse maintained at 27 ℃ (Lam et al. 2017). The 4-week-old rice seedlings (T_2_) for phenotypic and root cell wall analyses were grown in a growth chamber using a hydroponic system as described previously (Koshiba et al., 2013; Lam et al. 2024).

### Cell wall analyses

Extractive-free CWR samples were prepared from 5-month-old T_1_ (culm, leaf sheath, and leaf blade) or 4-week-old seedling T_2_ (root) mutant plants together with the WT control following the methods of Yamamura et al. (2012). Thioglycolic acid lignin assay (Suzuki et al., 2009), quantification of cell wall-bound FA and *p*CA by mild alkaline hydrolysis (Yamamura et al. 2011), analytical thioacidolysis (Yamamura et al., 2012; Yue et al., 2012), DFRC (Karlen et al., 2018; Takeda et al., 2018), and analysis of the polysaccharidic composition by two-step trifluoroacetic acid/sulfuric acid-catalyzed hydrolysis (Lam et al. 2017) were conducted as previously described. Histochemical analysis and 2D HSQC NMR were performed as described in **Supplemental Methods**. Peak assignments for HSQC plots are based on comparison with reported chemical shift data (Kim and Ralph 2010; Mansfield et al. 2012; Kim et al., 2017; Tobimatsu et al., 2019; Afifi et al. 2022; Martin et al., 2023).

### Isotopic feeding of ^13^C-labeled Phe and Tyr

Rice seeds were treated sequentially with 0.5% (v/v) hydrogen peroxide (overnight), 70% (v/v) ethanol (5 min), and 10% (v/v) sodium hypochlorite with one drop of TWEEN 20 (15 min) before germination. Ten seeds per experimental batch were germinated and grown in culture tubes containing ½MS medium (4 mL) supplemented with 3% (w/v) sucrose in 0.5% (w/v) Phytagel (pH 5.8) and 1 mM ^13^C_6_-Phe or ^13^C_9_-Tyr (Cambridge Isotope Laboratories) in a growth chamber maintained at 27 °C with a 12 h/12 h light/dark cycle, and harvested after 30 days for phenotypic and cell wall characterizations. Extractive-free CWR samples were prepared from the shoot parts via sequential extractions with water, 80% (v/v) ethanol, and acetone (Tobimatsu et al., 2013). Analytical thioacidolysis (Yamamura et al., 2012; Yue et al., 2012; Lan et al., 2016; Chen et al., 2021) and mild alkaline hydrolysis (Yamamura et al. 2011) followed by GC- or LC-MS analysis were used to determine the incorporation of ^13^C-labeled Phe and Tyr into lignin and hydroxycinnamate units (**H_thio_**, **G_thio_**, **S_thio_**, **T_thio_**, ***p*CA_AH_**, and **FA_AH_**), as described in **Supplemental Methods**.

### Chemical inhibitor assay

Surface-sterilized rice seeds were germinated on floating boats in tap water in a growth chamber maintained at 27 °C with a 12 h/12 h light/dark cycle. Three days after emergence, seedlings were transferred to new floating boat containers with a hydroponic solution (Koshiba et al., 2013) for two days before chemical inhibitor treatments. PA and AOPP were prepared as 50 mM stock solutions in 2 mL of DMSO. For plant chemical treatments, PA and AOPP stocks were diluted with distilled water to concentrations of 1 mM and 3 mM, respectively. Mock treatments consisted of 0.1% (v/v) DMSO diluted with distilled water. For each plant, approximately 1.5 mL of the diluted PA and AOPP solutions were added to 10 mL of hydroponic solution. All treated plants were grown in a growth chamber at 27 °C with a 12 h/12 h light/dark cycle for 9 days. Phenotypic performance was recorded by measuring plant height, culm length, and root length. The plants were then dried in an oven at 45 °C for one day. Thereafter, the aerial and root parts were weighed and subjected to CWR preparation and analytical thioacidolysis as described above.

### Statistical analysis

Student’s *t*-test and one way analysis of variance (ANOVA) followed by Tukey’s honestly significant difference test (*P* < 0.05) were performed using IBM SPSS Statistics version 27 (IBM Corporation, Armonk, NY, USA).

### Accession numbers

The accession numbers for the protein and gene sequences used in this study are listed in **Supplemental Table S1**.

## Supporting Information

**Supplemental Methods** Additional experimental procedures.

**Supplemental Figure S1.** Multiple alignment of C4H proteins from rice and other model plants.

**Supplemental Figure S2.** Predicted effects of CRISPR-Cas9-induced mutations on OsC4H1.

**Supplemental Figure S3.** Predicted effects of CRISPR-Cas9-induced mutations on OsC4H2a.

**Supplemental Figure S4.** Predicted effects of CRISPR-Cas9-induced mutations on OsC4H2b.

**Supplemental Figure S5.** 2D HSQC NMR spectra of *C4H*-knockout rice culm cell walls.

**Supplemental Figure S6.** Analysis of ^13^C-labeled lignin units by thioacidolysis.

**Supplemental Figure S7.** Analysis of ^13^C-labeled cell wall-bound hydroxycinnamates by alkaline hydrolysis.

**Supplemental Table S1.** Gene locus and accession numbers of C4H proteins examined.

**Supplemental Table S2.** Amino acid sequence identity of rice and Arabidopsis C4H proteins.

**Supplemental Table S3.** Oligonucleotide sequences used for constructing sgRNAs.

**Supplemental Table S4.** Off-target analyses of *C4H*-knockout rice mutants.

**Supplemental Table S5.** Neutral sugar analysis of *C4H*-knockout rice culm cell walls.

**Supplemental Table S6.** Peak assignments in 2D HSQC NMR spectra of rice cell walls.

**Supplemental Table S7.** Primers and oligonucleotides used in this study.

**Supplemental References**

## Acknowledgments

We thank Prof. Anne Britt and Dr. Mily Ron (University of California Davis) for providing the pMR285 CRISPR-Cas9 vector, Dr. Masaki Endo (National Agriculture and Food Research Organization) and Dr. Yuri Takeda-Kimura (Yamagata University) for helpful assistance in the rice genome editing experiments, and Prof. Hironori Kaji and Ms. Ayaka Maeno (ICR, Kyoto University) for their assistance in NMR analysis. A part of this study was conducted using the facilities at the DASH/FBAS, RISH, Kyoto University, and the NMR spectrometer at JURC, ICR, Kyoto University.

## Author contributions

S., L.P.Y.L., T.U. and Y.T. conceived research. S., L.P.Y.L., S.Y., O.A.A, P.J., Y.O., K.O. and Y.T.. designed and performed experiments, and analyzed the data. S., L.P.Y.L., S.Y., P.J. and Y.T. wrote the manuscript with help from all the others.

## Data availability

The authors confirm that the data supporting the findings of this study are available within the article and its supplementary materials.

