## Supplemental Materials for "Essential Yet Dispensable: The Role of CINNAMATE 4-HYDROXYLASE in Rice Cell Wall Lignification"

[a]Research Institute for Sustainable Humanosphere, Kyoto University, Gokasho, Uji, Kyoto 611-0011, Japan; [b]Center for Crossover Education, Graduate School of Engineering Science, Akita University, Tegata Gakuen-machi 1-1, Akita City, Akita 010-8502, Japan; [c]School of Life Science and Technology, Tokyo Institute of Technology, Kanagawa 226-8502 Japan; [d]Faculty of Bioscience and Bioindustry, Tokushima University, Tokushima 770-8503 Japan.

[†]These authors contributed equally to this study.

[‡]Present address: Biology Department, Brookhaven National Laboratory, Upton, NY, USA.

**List of Materials**

**Supplemental Methods** Additional experimental procedures.

**Supplemental Figure S1.** Multiple alignment of C4H proteins from rice and other model plants.

**Supplemental Figure S2.** Predicted effects of CRISPR/Cas9-induced mutations on OsC4H1.

**Supplemental Figure S3.** Predicted effects of CRISPR/Cas9-induced mutations on OsC4H2a.

**Supplemental Figure S4.** Predicted effects of CRISPR/Cas9-induced mutations on OsC4H2b.

**Supplemental Figure S5.** 2D HSQC NMR spectra of *C4H*-knockout rice culm cell walls.

**Supplemental Figure S6.** Analysis of $^{13}C$-labeled lignin units by thioacidolysis.

**Supplemental Figure S7.** Analysis of $^{13}C$-labeled cell wall-bound hydroxycinnamates by alkaline hydrolysis.

**Supplemental Table S1.** Gene locus and accession numbers of C4H homologs examined.

**Supplemental Table S2.** Amino acid sequence identity of rice C4H members.

**Supplemental Table S3.** Oligonucleotide sequences used for constructing sgRNAs.

**Supplemental Table S4.** Off-target analyses of *C4H*-knockout rice mutants.

**Supplemental Table S5.** Neutral sugar analysis of *C4H*-knockout rice culm cell walls.

**Supplemental Table S6.** Peak assignments in 2D HSQC NMR spectra of rice cell walls.

**Supplemental Table S7.** Primers and oligonucleotides used in this study.

**Supplemental References**

### Supplemental Methods

*Plant transformation and growth conditions*

To generate *OsC4H1*-KO and *OsC4H2a/2b*-DKO, the sgRNAs targeting *OsC4H1* and conserved regions of *OsC4H2a* and *OsC4H2b* were designed with minimum off-target potential using CRISPR-P 2.0 (**Supplemental Table S3**) (Liu et al. 2017), and cloned into pMR285 binary vector (Ritter *et al.*, 2017) using the oligonucleotides listed in **Supplemental Table S7**. The obtained sgRNA-Cas9 binary vectors were transformed into embryogenic calli derived from wild-type rice (cv. Nipponbare) seeds via *Agrobacterium tumefaciens* strain EHA101 and regenerated following the methods of Hiei et al. (1994) and Hattori et al. (2012). To generate *OsC4H1/2a/2b*-TKO, a homozygous *OsC4H2a/2b*-DKO line without the sgRNA-Cas9 cassette was first isolated (as described below) and subsequently transformed with the sgRNA-Cas9 vector targeting *OsC4H1*. The regenerated ($T_0$) plants were genotyped and grown to maturity in potting soil in a growth chamber (Lam et al. 2017). Selected $T_1$ and $T_2$ mutant lines were further genotyped and grown to maturity in a greenhouse maintained at 27 °C (Lam et al. 2017). The 4-week-old rice seedlings ($T_2$) for phenotypic and root cell wall analyses were grown in a growth chamber using a hydroponic system as described previously (Lam et al. 2024). For genotyping, genomic DNA was extracted from young leaves and the genomic region containing the target site was amplified by PCR using primers listed in **Supplemental Table S7**. The amplified PCR products were subjected to direct sequencing in accordance with the method of Takeda et al. (2019). Potential off-target sites were predicted with CRISPR-P 2.0 (**Supplemental Table S4**) (Liu et al. 2017). Genomic regions, including the three top-ranked potential off-target sites, were amplified by PCR using primers listed in **Supplemental Table S7** and subjected to direct sequencing as described above.

*Histochemical analysis*

Specimens (approximately 1 cm) were manually excised from culms at the heading stage and then fixed in a formaldehyde/propionic acid/ethanol (3.5:5:50, v/v/v) solution and decolorized in an ethanol/acetic acid (6:1, v/v) solution. Hand-cut sections were subjected to lignin staining with 1% (w/v) phloroglucinol in ethanol for 10 min, acidified with 17.5% (w/w) HCl, and incubated for an additional 10 min. The stained sections were then examined using an Olympus BX51 microscope (Olympus Optical, Tokyo, Japan).

*2D HSQC NMR*

For 2D HSQC NMR analysis, the culm and leaf sheath cell wall residue (CWR) samples (~60 mg) were ball-milled (Afifi et al. 2022) and swollen in 600 µl solvent mixture of dimethyl sulfoxide-$d_6$ (DMSO-$d_6$)/pyridine-$d_5$ (4:1, v/v) for gel-state NMR analysis as described previously (Kim and Ralph 2010; Mansfield et al. 2012). The HSQC NMR spectra were acquired using a Bruker Biospin Avance III 800 system (800 MHz, Bruker Biospin, Billerica, MA, USA) equipped with a cryogenically cooled 5 mm TCI gradient probe. Adiabatic HSQC NMR experiments were conducted using the standard Bruker implementation ("hsqcetgpsp.3"). Specific parameters were set as described previously (Kim and Ralph 2010; Mansfield et al. 2012). Data processing and analysis were performed as described previously (Afifi et al. 2022; Martin et al., 2023) using Bruker TopSpin 4.0 (Mac) software (Bruker Biospin). The central DMSO solvent peaks ($\delta_C/\delta_H$: 39.5/2.49 ppm) were used as internal references. Peak assignments for HSQC plots are listed in **Supplemental Table S6** (Kim and Ralph 2010; Mansfield et al. 2012; Kim et al., 2017; Tobimatsu et al., 2019; Afifi et al. 2022; Martin et al., 2023). For volume integration analysis of HSQC plots, the well-resolved aromatic C2–H2 correlations from lignin ($\mathbf{G_2}$ and $\mathbf{S_{2/6}}$), hydroxycinnamates ($\boldsymbol{p}\mathbf{CA_{2/6}}$ and $\mathbf{FA_2}$), and tricin ($\mathbf{T_{2'/6'}}$) units, and anomeric C1–H1 correlations from polysaccharide units ($\mathbf{Gl_1}$, $\mathbf{Xy_1}$, $\mathbf{Xy'_1}$, $\mathbf{Xy''_1}$, $\mathbf{Ar_1}$, $\mathbf{Ga_1}$, and $\mathbf{GlU_1}$) were manually integrated and signals from $\mathbf{H_{2/6}}$, $\mathbf{S_{2/6}}$, $\mathbf{P_{2/6}}$, and $\mathbf{T_{2'/6'}}$ were logically halved (Afifi et al. 2022). Each signal was normalized according to the sum of the integrated signals ($\frac{1}{2}\mathbf{S_{2/6}} + \mathbf{G_2} + \frac{1}{2}\mathbf{T_{2'/6'}} + \mathbf{FA_2} + \frac{1}{2}\mathbf{P_{2/6}} + \mathbf{Gl_1} + \mathbf{Xy_1} + \mathbf{Xy'_1} + \mathbf{Xy''_1} + \mathbf{Ar_1} + \mathbf{Ga_1} + \mathbf{GlU_1} = 100$). For volume integration analysis of lignin inter-monomeric linkages, the well-resolved Cα–Cα contours from β-*O*-4 ($\mathbf{I_\alpha}$), β-5 ($\mathbf{II_\alpha}$), and β-β ($\mathbf{III'_\alpha}$) units were used, and the $\mathbf{III'_\alpha}$ integral was logically halved.

*Isotopic feeding of $^{13}$C-labeled Phe and Tyr*

Extractive-free CWR samples prepared from rice seedlings fed with $^{3}C_6$-Phe or $^{13}C_9$-Tyr (see **Materials and Methods** in the main article) were subjected analytical thioacidolysis and mild alkaline hydrolysis to determine the incorporation of $^{13}$C-labeled Phe and Tyr into lignin and hydroxycinnamate units.

Analytical thioacidolysis was performed following the previously described methods (Yamamura et al., 2012; Yue et al., 2012; Lan et al., 2016; Chen et al., 2021) with some modifications. Briefly, CWR (~5 mg) was treated in a freshly prepared thioacidolysis reagent (dioxane/ethanethiol/$BF_3$-etherate, 35:4:1, v/v/v) at 100 °C for 4 h, cooled down on ice, and then added with 4,4′-ethylidenebisphenol (Yue et al., 2012) for the analysis of monolignol-derived lignin units ($\mathbf{H_{thio}}$, $\mathbf{G_{thio}}$ and $\mathbf{S_{thio}}$), or umbelliferone (Lan et al., 2016) for the analysis ticin-lignin units ($\mathbf{T_{thio}}$) as an internal standard. The reaction mixture was then neutralized with 100 µL of 1 M $NaHCO_3$ solution, and extracted three times with ~250 µL ethyl acetate. The combined organic layer was washed three times with saturated NaCl, dried over $Na_2SO_4$, and concentrated *in vacuo*. For the analysis of monolignol-derived lignin units (**$H_{thio}$**, **$G_{thio}$**, and **$S_{thio}$**), an aliquot of the thioacidolysis reaction mixture was derivatized in *N*,*O*-bis(trimethylsilyl)-acetamide (BSA) and subjected to GC-MS on a Shimadzu QP-2010 Plus (Shimadzu, Kyoto, Japan) under the following conditions: column, Shimadzu HiCap CBP10 (25 m × 0.22 mm); carrier gas, helium; injection temperature, 230 °C; oven temperature, 40 °C at t = 0 to 2 min, then to 230 °C at 40 °C $min^{-1}$ elevation rate; ion source, 300 °C; interface temperature, 250 °C; ionization, electron-impact mode (70 eV); MS detection by selective ion mode (SIM), *m/z* = 239, 245, and 246 for unlabeled, $^{13}C_6$-labeled and $^{13}C_9$-labeled **$H_{thio}$**, *m/z* = 269, 275, and 276 for unlabeled, $^{13}C_6$-labeled and $^{13}C_9$-labeled **$G_{thio}$**, *m/z* = 299, 305, and 306 for unlabeled, $^{13}C_6$-labeled and $^{13}C_9$-labeled **$S_{thio}$**, *m/z* = 343 for 4,4′-ethylidenebisphenol (internal standard). For the analysis of tricin-lignin unit (**$T_{thio}$**) (Lan et al., 2016; Chen et al., 2021), an aliquot of the thioacidolysis reaction mixture was resuspended in 200 μL of 95% (v/v) methanol, filtered through a 0.45 μm membrane filter (Cosmonice Filter S, Nacalai Tesque, Kyoto, Japan), and subjected to LC-MS on a Shimadzu LCMS-2020 (Shimadzu, Kyoto, Japan) with the following conditions: injection volume, 10 μL; column, Kinetex 5 μl XB-C18 100 Å LC Column 250 × 4.6 mm (Phenomenex, Chicago, IL, USA); eluent, 60% (v/v) methanol with 0.1% (v/v) formic acid; flow rate, 0.4 ml/min; MS desolvation line, 250 ℃; heat block, 200 ℃; MS nebulizer gas, $N_2$ at 1.5 $min^{-1}$; MS ionization, ESI (positive ion mode); MS detection by selective ion mode (SIM), *m/z* = 331, 337, and 340 for unlabeled, $^{13}C_6$-labeled, and $^{13}C_9$-labeled **$T_{thio}$** ($[M+H]^+$), and *m/z* = 163 for umbelliferone (internal standard) ($[M+H]^+$).

Mild alkaline hydrolysis was performed according to Yamamura et al. (2011) with minor modifications. Briefly, CWR (~5 mg) was treated in 1 M NaOH (1.5 mL) with gentle shaking at 25 ℃ for 24 h in the dark. After centrifugation (16,100 g, 10 min at room temperature), the supernatant was added with *o*-coumaric acid as an internal standard. The reaction mixture was then neutralized with 1.2 mL of 2N HCl solution, and extracted three times with ~250 μL ethyl acetate. The combined organic layer was washed three times with saturated NaCl, dried over $Na_2SO_4$, and concentrated *in vacuo*. An aliquot of the reaction mixture of mild alkaline hydrolysis was derivatized in BSA and subjected to GC-MS on a Shimadzu QP-2010 Plus (Shimadzu, Kyoto, Japan) under the following conditions: column, Shimadzu HiCap CBP10 (25 m × 0.22 mm); carrier gas, helium; injection temperature, 230 °C; oven temperature, 40 °C at t = 0 to 2 min, then to 230 °C at 40 °C $min^{-1}$ elevation rate; ion source temperature, 300 °C; interface temperature, 250 °C; ionization, electron-impact mode (70 eV); MS detection by selective ion mode (SIM), *m/z* = 308, 314, and 317 for unlabeled, $^{13}C_6$-labeled, and $^{13}C_9$-labeled ***p*CA$_{AH}$**, *m/z* = 338, 344, and 347 for unlabeled, $^{13}C_6$-labeled, and $^{13}C_9$-labeled **$FA_{AH}$**, *m/z* = 308 for *o*-coumaric acid (internal standard).

The levels of $^{13}C$-labeled products were estimated by the MS peak areas of the labeled and unlabeled thioacidolysis products using the following equations (Barros et al., 2016):

$$\%^{13}\text{C incorporation} = \frac{\text{peak area of } ^{13}\text{C-labeled ion}}{(\text{peak area of } ^{13}\text{C-labeled ion} + \text{peak area of unlabeled ion})} \times 100$$

$$\%^{13}\text{C-Phe/Tyr proportion} = \frac{^{13}\text{C-Phe/Tyr incorporation}}{(^{13}\text{C-Phe incorporation} + {}^{13}\text{C-Tyr incorporation})} \times 100$$

| Class | Sequence | Alignment | Position |
|---|---|---|---|
| class I | ***Oryza sativa* OsC4H1b (CYP73A38)** | -----------------------MDALL----------VEKVLLGLFVAAVLALVVAKLT | 27 |
|  | ***Oryza sativa* CYP73A35p** | -----------------------MDLLF----------VERLLVGLLAAAVVAIAVSKLR | 27 |
|  | *Brachypodium distachyon* Bradi2g31510 | -----------------------MDVLL----------LEKALLGLFAAAVLAIAVAKLT | 27 |
|  | *Brachypodium distachyon* Bradi2g53470 | -----------------------MDFLF----------VEKLLVGLLASALVAIAVSKIR | 27 |
|  | *Arabidopsis thaliana* AtC4H | -----------------------MDLLL----------LEKSLIAVFVAVILATVISKLR | 27 |
|  | *Populus trichocarpa* Potri013G157900 | -----------------------MDLLL----------LEKTLLGSFVAILVAILVSKLR | 27 |
|  | *Populus trichocarpa* Potri019G130700 | -----------------------MDLLL----------LEKTLLGSFVAILVAILVSQLR | 27 |
| class II | ***Oryza sativa* OsC4H2a (CYP73A39)** | MAASVVRV---AIAT-GAS--LA-VHLFVKSFLQAQHPALTLLLPVAVFAGIAVGAKGGN | 53 |
| | ***Oryza sativa* OsC4H2b (CYP73A40)** | MAASAMRV---AIAT-GAS--LA-VHLFVKSFVQAQHPALTLLLPVAVFVGIAVGAKGGS | 53 |
| | *Brachypodium distachyon* Bradi3g43160 | MAALAIRA---AFAAVATS--LAVYWLLNSSFLQTPNIALSLPAAAAAFVVVAIAASGPG | 55 |
| | *Populus trichocarpa* Potri018G146100 | MASFVTKSMGFTLLAVASVSCIKFACPNLSTYFSPL--PISVILPLLPLIVYLFSSVFTK | 58 |

**cluster of proline**

| Class | Sequence | Alignment | Position |
|---|---|---|---|
| class I | ***Oryza sativa* OsC4H1b (CYP73A38)** | GKRLRL**PPGPAGAP**IVGNWLQVGDDLNHRNLMALARRFGDILLLRMGVRNLVVVSSPDLA | 87 |
| | ***Oryza sativa* CYP73A35p** | GRKLRL**PPGPTPVP**VFGNWLQVGDDLNHRNLAALARRFGDIFLLRMGQRNLVVVSSPPLA | 87 |
| | *Brachypodium distachyon* Bradi2g31510 | GKRFRL**PPGPSGAP**IVGNWLQVGDDLNHRNLMGMAKRFGEVFLLRMGVRNLVVVSSPDLA | 87 |
| | *Brachypodium distachyon* Bradi2g53470 | GRKLKL**PPGPLPVP**IFGNWLQVGDDLNHRNLAAMARKFGEIFLLRMGQRNLVVVSSPPLA | 87 |
| | *Arabidopsis thaliana* AtC4H | GKKLKL**PPGPIPIP**IFGNWLQVGDDLNHRNLVDYAKKFGDLFLLRMGQRNLVVVSSPDLT | 87 |
| | *Populus trichocarpa* Potri013G157900 | GKRFKL**PPGPIPVP**VFGNWLQVGDDLNHRNLTDLAKKFGDIFLLRMGQRNLVVVSSPDLS | 87 |
| | *Populus trichocarpa* Potri019G130700 | GKRFKL**PPGPLPVP**VFGNWLQVGDDLNHRNLTDLAKKFGDILLLRMGQRNLVVVSSPDLA | 87 |
| class II | ***Oryza sativa* OsC4H2a (CYP73A39)** | GGDGKA**PPGPAAVP**VFGNWLHVGNDLNHRFLAAMSARYGPVFRLRLGVRNLVVVSDPKLA | 113 |
| | ***Oryza sativa* OsC4H2b (CYP73A40)** | GGDGKA**PPGPAAVP**VFGNWLQVGNDLNHRFLAAMSARYGPVFRLRLGVRNLVVVSDPKLA | 113 |
| | *Brachypodium distachyon* Bradi3g43160 | HRSDGT**PPGPAALP**VLGNWLQVGNDLNHRFLARLSARYGPVFRLRLGVRNLVVVSDPRLA | 115 |
| | *Populus trichocarpa* Potri018G146100 | SSTGDL**PPGPVSYP**MFGNWLQVGNDLNHRLLASMSQTYGPVFLLKLGSKNLAVVSDPELA | 118 |

| Class | Sequence | Alignment | Position |
|---|---|---|---|
| class I | ***Oryza sativa* OsC4H1b (CYP73A38)** | KEVLHTQGVEFGSRTRNVVFDIFTGKGQDMVFTVYGDHWRKMRRIMTVPFFTNKVVAQNR | 147 |
| | ***Oryza sativa* CYP73A35p** | REVLHTQGVEFGSRTRNVVFDIFTGKGQDMVFTVYGDHWRKMRRIMTVPFFTGKVVQRHR | 147 |
| | *Brachypodium distachyon* Bradi2g31510 | KEVLHTQGVEFGSRTRNVVFDIFTGKGQDMVFTVYGDHWRKMRRIMTVPFFTNKVVAQNR | 147 |
| | *Brachypodium distachyon* Bradi2g53470 | REVLHTQGVEFGSRTRNVVFDIFTGEGQDMVFTVYGDHWRKMRRIMTVPFFTNKVVQQYR | 147 |
| | *Arabidopsis thaliana* AtC4H | KEVLLTQGVEFGSRTRNVVFDIFTGKGQDMVFTVYGEHWRKMRRIMTVPFFTNKVVQQNR | 147 |
| | *Populus trichocarpa* Potri013G157900 | KEVLHTQGVEFGSRTRNVVFDIFTGKGQDMVFTVYGEHWRKMRRIMTVPFFTNKVVQQYR | 147 |
| | *Populus trichocarpa* Potri019G130700 | KEVLHTQGVEFGSRTRNVVFDIFTGKGQDMVFTVYGEHWRKMRRIMTVPFFTNKVVQQYR | 147 |
| class II | ***Oryza sativa* OsC4H2a (CYP73A39)** | TEVLHTQGVEFGSRPRNVVFDIFTANGADMVFTEYGDHWRRMRRVMTLPFFTARVVQQYK | 173 |
| | ***Oryza sativa* OsC4H2b (CYP73A40)** | TEVLHTQGVEFGSRPRNVVFDIFTANGADMVFTEYGDHWRRMRRVMTLPFFTARVVQQYK | 173 |
| | *Brachypodium distachyon* Bradi3g43160 | TEVLHTQGVEFGSRPRNVVFDIFTANGADMVFTEYGDHWRRMRRVMTLPFFTARVVQQYR | 175 |
| | *Populus trichocarpa* Potri018G146100 | NQVLHTQGVEFGSRPRNVVFDIFTGNGQDMVFTIYGEHWRKMRRIMTLPFFTNKVVNQYS | 178 |

**truncation in CYP73A35p**

| Class | Sequence | Alignment | Position |
|---|---|---|---|
| class I | ***Oryza sativa* OsC4H1b (CYP73A38)** | AGWEEEARL**VVEDVRRDPAAATSGVVIRRRL**QLMMYNDMFRIMFDRRFDSVDDPLFNKLK | 207 |
|  | ***Oryza sativa* CYP73A35p** | AGWEAEAAA----------------------VLMMYSNVYRIMFDRRFESADDPLFLRLK | 185 |
| | *Brachypodium distachyon* Bradi2g31510 | VGWEEEARL**VVEDVRADPASAVGGVVIRRRL**QLMMYNDMFRIMFDRRFASVDDPLFNKLK | 207 |
| | *Brachypodium distachyon* Bradi2g53470 | AGWEAEAAF**VVDNVRADPRAATDGVVLRRHL**QLMMYNNMYRIMFDRRFESLDDPLFLRLR | 207 |
| | *Arabidopsis thaliana* AtC4H | EGWEFEAAS**VVEDVKKNPDSATKGIVLRKRL**QLMMYNNMFRIMFDRRFESEDDPLFLRLK | 207 |
| | *Populus trichocarpa* Potri013G157900 | YGWEEEAAQ**VVEDVKKNPEAATNGIVLRRRL**QLMMYNNMYRIMFDRRFESEDDPLFNKLK | 207 |
| | *Populus trichocarpa* otri019G130700 | YGWEEEAAQ**VVEDVKKNPEAATHGIVLRRRL**QLMMYNNMYRIMFDRRFESEEDPLFNKLK | 207 |
| class II | ***Oryza sativa* OsC4H2a (CYP73A39)** | AMWEAEMDA**VVDDVRGDAVAQGTGFVVRRRL**QLMLYNIMYRMMFDARFESVDDPMFIEAT | 233 |
| | ***Oryza sativa* OsC4H2b (CYP73A40)** | AMWEAEMDA**VVDDVRGDAVAQGTGFVVRRRL**QLMLYNIMYRMMFDARFESVDDPMFIEAT | 233 |
| | *Brachypodium distachyon* Bradi3g43160 | AMWEAEMDA**VVSDLRADPVARVAGVVVRRRL**QLMLYNIMYGMMFDARFESVDDPLFVQAT | 235 |
| | *Populus trichocarpa* Potri018G146100 | TSWEQEMDL**VVDDLRANEKVRTEGIVIRKRL**QLMLYNIMYRMMFDAKFQSQEDPLFVQAT | 238 |

| Class | Sequence | Alignment | Position |
|---|---|---|---|
| class I | ***Oryza sativa* OsC4H1b (CYP73A38)** | AFNAERSRLSQSFEYNYGDFIPVLRPFLRRYLARCHQLKSQRMKLFEDHFVQERKRVME- | 266 |
| | ***Oryza sativa* CYP73A35p** | ALNGERSRLAQSFEYNYGDFIPILRPFLRGYLRICEEVKETRLKLFKDFFLEERKKLAST | 245 |
| | *Brachypodium distachyon* Bradi2g31510 | ALNAERSILSQSFDYNYGDFIPVLRPFLRRYLNRCHTLKSKRMKVFEDHFVQERKEALE- | 266 |
| | *Brachypodium distachyon* Bradi2g53470 | ALNGERSRLAQSFDYNYGDFIPILRPFLRGYLRLCKEVKETRLKLFKDYFLEERKKLAST | 267 |
| | *Arabidopsis thaliana* AtC4H | ALNGERSRLAQSFEYNYGDFIPILRPFLRGYLKICQDVKDRRIALFKKYFVDERKQIASS | 267 |
| | *Populus trichocarpa* Potri013G157900 | ALNGERSRLAQSFDYNYGDFIPILRPFLRGYLKICQEVKERRLQLFKDYFVDERKKLAST | 267 |
| | *Populus trichocarpa* Potri019G130700 | ALNGERSRLAQSFDYNYGDFIPILRPFLRGYLKICKEVKERRLQLFKDYFVEERKKLGST | 267 |
| class II | ***Oryza sativa* OsC4H2a (CYP73A39)** | RFNSERSRLAQSFEYNYGDFIPILRPFLRGYLNKCRDLQSRRLAFFNNNYVEKRRKVMDT | 293 |
| | ***Oryza sativa* OsC4H2b (CYP73A40)** | RFNSERSRLAQSFEYNYGDFIPILRPFLRGYLNKCRDLQSRRLAFFNNNYVEKRRKVMDT | 293 |
| | *Brachypodium distachyon* Bradi3g43160 | RFNSERSRLAQSFDYNYGDFIPILRPFLRGYLNKCRDLQSRRLAFFNNNYVEKRRKVMDS | 295 |
| | *Populus trichocarpa* Potri018G146100 | RFNSERSRLAQSFEYNYGDFIPWLRPFLRGYLNKCRDLQQRRLAFFNNYYIEKRRKIMAA | 298 |

**oxygen binding region**

| Class | Sequence | Alignment | Position |
|---|---|---|---|
| class I | ***Oryza sativa* OsC4H1b (CYP73A38)** | ---QTGEIRCAMDHILEAERKGEINHDNVLYIVENINV**AAIETT**LWSIEWGIAELVNHPS | 323 |
| | ***Oryza sativa* CYP73A35p** | KAMDNNGLKCAIDHILEAQQKGEINEDNVLYIVENINV**ADDAVV--**DGVGDRGARVNHGE | 303 |
| | *Brachypodium distachyon* Bradi2g31510 | ---KTGEIRCAMDHILEAERKGEINHDNVLYIVENINV**AAIETT**LWSIEWGIAELVNHPE | 323 |
| | *Brachypodium distachyon* Bradi2g53470 | KAMDNNGLKCAIDHILEAQQKGEINEDNVLYIIENINV**AAIETT**LWSIEWAIAELVNHPE | 327 |
| | *Arabidopsis thaliana* AtC4H | KPTGSEGLKCAIDHILEAEQKGEINEDNVLYIVENINV**AAIETT**LWSIEWGIAELVNHPE | 327 |
| | *Populus trichocarpa* Potri013G157900 | KNMNNEGLKCAIDHILDAQKKGEINEDNVLYIVENINV**AAIETT**LWSIEWGIAELVNHPE | 327 |
| | *Populus trichocarpa* Potri019G130700 | KSMSNEGLKCAIDHILDAQKKGEINEDNVLYIVENINV**AAIETT**LWSIEWGIAELVNHPE | 327 |
| class II | ***Oryza sativa* OsC4H2a (CYP73A39)** | P-GDRNKLRCAIDHILEAEKNGELTAENVIYIVENINV**AAIETT**LWSIEWALAEVVNHPA | 352 |
| | ***Oryza sativa* OsC4H2b (CYP73A40)** | P-GDRNKLRCAIDHILEAEKNGELTAENVIYIVENINV**AAIETT**LWSIEWALAEVVNHPA | 352 |
| | *Brachypodium distachyon* Bradi3g43160 | P-GDKDKLRCAIDHILAAEKNGEITAENVIYIVENINV**AAIETT**LWSIEWALAEVVNHPA | 354 |
| | *Populus trichocarpa* Potri018G146100 | N-GEKHKVSCAMDHIIQAQMKGEISEENVLYIVENINV**AAIETT**LWSMEWAIAELVNHPT | 357 |

| Class | Protein | Sequence | Position |
|---|---|---|---|
| class I | *Oryza sativa* OsC4H1b (CYP73A38) | IQSKVREEMASVLGG-AAVTEPDLERLPYLQAVVKETLRLRMAIPLLVPHMNLADGKLAG | 382 |
| | *Oryza sativa* CYP73A35p | IQEKLRRELDTVLGPGRQITEPDTHRLPYLQAVVKETLRLRMAIPLLVPHMNLRDAELAG | 363 |
| | *Brachypodium distachyon* Bradi2g31510 | IQAKVREEITAVLGPNTAVTEPDLERLPYLQAVVKETLRLRMAIPLLVPHMNLSDAKLAG | 383 |
| | *Brachypodium distachyon* Bradi2g53470 | IQQKLRDELDTVLGAGHQITEPDTHKLPYLQAVIKETLRLRMAIPLLVPHMNLQDAKLGG | 387 |
| | *Arabidopsis thaliana* AtC4H | IQSKLRNELDTVLGPGVQVTEPDLHKLPYLQAVVKETLRLRMAIPLLVPHMNLHDAKLAG | 387 |
| | *Populus trichocarpa* Potri013G157900 | IQKKLRHELDTLLGPGHQITEPDTYKLPYLNAVIKETLRLRMAIPLLVPHMNLHDAKLGG | 387 |
| | *Populus trichocarpa* Potri019G130700 | IQKKLRDELDTVLGPGHQITEPDTNKLPYLNAVIKETLRLRMAIPLLVPHMNLHDAKLGG | 387 |
| class II | *Oryza sativa* OsC4H2a (CYP73A39) | VQSKVRAEINDVLGDDEPITESSIHKLTYLQAVIKETLRLHSPIPLLVPHMNLEEAKLGG | 412 |
| | *Oryza sativa* OsC4H2b (CYP73A40) | VQSKVRAEINDVLGDDEPITESSIHKLTYLQAVIKETLRLHSPIPLLVPHMNLEEAKLGG | 412 |
| | *Brachypodium distachyon* Bradi3g43160 | VQTKVRGEIKDVLGDDEPITESNIQQLPYLQAVIKETLRLHSPIPLLVPHMNLEEAKLGG | 414 |
| | *Populus trichocarpa* Potri018G146100 | VQRKIRDEIRAVLK-GSPVTESNLHELPYLQATIKETLRLHTPIPLLVPHMNLEEAKLGG | 416 |

heme binding region

| Class | Protein | Sequence | Position |
|---|---|---|---|
| class I | *Oryza sativa* OsC4H1b (CYP73A38) | YDIPAESKILVNAWFLANDPKRWVRPDEFRPERFLEEEKAVEA----HGNDFRFVP**FGVG** | 438 |
|  | *Oryza sativa* CYP73A35p | YGIPAESKVLVNAWYLANDPGRWRRPEEFRPERFLEEERNVEA----NGNDFRYLP**SGAG** | 419 |
|  | *Brachypodium distachyon* Bradi2g31510 | YDIPAESKILVNAWFLANDPKRWVRPDEFRPERFMEEEKAVEA----HGNDFRFVP**FGVG** | 439 |
|  | *Brachypodium distachyon* Bradi2g53470 | YNIPAESKILVNAWFLANNPEEWRRPDEFRPERFLEEEKHVEA----NGNDFRFLP**FGVG** | 443 |
|  | *Arabidopsis thaliana* AtC4H | YDIPAESKILVNAWWLANNPNSWKKPEEFRPERFFEEESHVEA----NGNDFRYVP**FGVG** | 443 |
|  | *Populus trichocarpa* Potri013G157900 | FDIPAESKILVNAWWLANNPAHWKNPEEFRPERFLEEEAKVEA----NGNDFRYLP**FGVG** | 443 |
|  | *Populus trichocarpa* Potri019G130700 | FDIPAESKILVNAWWLANNPAKWKNPEEFRPERFFEEEAKVEA----NGNDFRYLP**FGVG** | 443 |
| class II | *Oryza sativa* OsC4H2a (CYP73A39) | YTIPKGSKVVVNAWWLANNPALWENPEEFRPERFLEKESGVDA-TVAGKVDFRFLP**FGVG** | 471 |
| | *Oryza sativa* OsC4H2b (CYP73A40) | YTIPKGSKVVVNAWWLANNPALWENPEEFRPERFLEKESGVDA-TVAGKVDFRFLP**FGVG** | 471 |
| | *Brachypodium distachyon* Bradi3g43160 | YTIPRGSKVVVNAWWLANNPELWEKPEEFRPERFLDEDSGVDAATIGGKADFRFLP**FGVG** | 474 |
| | *Populus trichocarpa* Potri018G146100 | FTIPKESKVVVNAWWLANNPEWWEKPSEFRPERFLEEERDTEA-IVGGKVDFRFLP**FGVG** | 475 |

heme binding region

| Class | Protein | Sequence | Position |
|---|---|---|---|
| class I | *Oryza sativa* OsC4H1b (CYP73A38) | **RRSCPG**IILALPIIGITLGRLVQSFDLLPPPGMDKVDTTEKPGQFSNQILKHATVVCKPI | 498 |
| | *Oryza sativa* CYP73A35p | **RRSCPG**IVLALPILGVTIGRLVQNFELLPPPGKDRVDTTEKGGQFSLHILKHSTIVAKPR | 479 |
| | *Brachypodium distachyon* Bradi2g31510 | **RRSCPG**IILALPIIGITLGRLVQNFTLLPPPGQAKIDTTEKPGQFSNQILKHATVVCKPL | 499 |
| | *Brachypodium distachyon* Bradi2g53470 | **RRSCPG**IILALPILGITIGRLVQNFELLPPPGQDKLDTTEKGGQFSLHILKHSNIVAKPR | 503 |
| | *Arabidopsis thaliana* AtC4H | **RRSCPG**IILALPILGITIGRMVQNFELLPPPGQSKVDTSEKGGQFSLHILNHSIIVMKPR | 503 |
| | *Populus trichocarpa* Potri013G157900 | **RRSCPG**IILALPILGITLGRLVQNFELLPPPGQSKIDTAEKGGQFSLHILKHSTIVAKPR | 503 |
| | *Populus trichocarpa* Potri019G130700 | **RRSCPG**IILALPILGITLGRLVQNFELLPPPGQSKIDTSEKGGQFSLHILKHSTIVAKPR | 503 |
| class II | *Oryza sativa* OsC4H2a (CYP73A39) | **RRSCPG**IILALPILALIVGKLVRSFEMVPPPGVEKLDVSEKGGQFSLHIAKHSVVAFHPI | 531 |
| | *Oryza sativa* OsC4H2b (CYP73A40) | **RRSCPG**IILALPILALIVGKLVRSFEMVPPPGVEKLDVSEKGGQFSLHIAKHSVVAFHPI | 531 |
| | *Brachypodium distachyon* Bradi3g43160 | **RRSCPG**IILAMPILALIVGKLVRSFQMLPPPGVDKLDVSEKGGQFSLHIANHSVVAFHPI | 534 |
| | *Populus trichocarpa* Potri018G146100 | **RRSCPG**IILAMPILGLIVARLVSNFEMIAPPGMEKIDVSEKGGQFSLHIASHSTVVFKPI | 535 |

| Class | Protein | Sequence | Position |
|---|---|---|---|
| class I | *Oryza sativa* OsC4H1b (CYP73A38) | DA- | 500 |
| | *Oryza sativa* CYP73A35p | AF- | 481 |
| | *Brachypodium distachyon* Bradi2g31510 | QA- | 501 |
| | *Brachypodium distachyon* Bradi2g53470 | VF- | 505 |
| | *Arabidopsis thaliana* AtC4H | NC- | 505 |
| | *Populus trichocarpa* Potri013G157900 | SF- | 505 |
| | *Populus trichocarpa* Potri019G130700 | SF- | 505 |
| class II | *Oryza sativa* OsC4H2a (CYP73A39) | SA- | 533 |
| | *Oryza sativa* OsC4H2b (CYP73A40) | SA- | 533 |
| | *Brachypodium distachyon* Bradi3g43160 | DSA | 537 |
| | *Populus trichocarpa* Potri018G146100 | KA- | 537 |

**Supplemental Figure S1. Multiple alignment of C4H proteins from rice and other model plants.** Conserved cytochrome P450 motifs (proline cluster, and oxygen and heme binding domains) (Chapple, 1998) and regions where CYP73A35P has a ~20 bp deletion are highlighted.

>OsC4H1

```
                                                                   cluster of proline
WT                         MDALLVEKVLLGLFVAAVLALVVAKLTGKRLRLPPGPAGAPIVGNWLQVGDDLNHRNLMA     60
OsC4H1-KO                  MDALLVEKVLLGLFVAAVLALVVAKLTGKRLRLPPGPAGAPIVGNWLQVGDDLNHRNLMA     60
OsC4H1/2a/2b-TKO-a         MDALLVEKVLLGLFVAAVLALVVAKLTGKRLRLPPGPAGAPIVGNWLQVGDDLNHRNLMA     60
OsC4H1/2a/2b-TKO-b         MDALLVEKVLLGLFVAAVLALVVAKLTGKRLRLPPGPAGAPIVGNWLQVGDDLNHRNLMA     60

WT                         LARRFGDILLLRMGVRNLVVVSSPDLAKEVLHTQGVEFGSRTRNVVFDIFTGKGQDMVFT    120
OsC4H1-KO                  LARRFGDILLLRMGVRNLVVVSSPDLAKEVLHTQGVEFGSRTRNVVFDIFTGKGQDMVFT    120
OsC4H1/2a/2b-TKO-a         LARRFGDILLLRMGVRNLVVVSSPDLAKEVLHTQGVEFGSRTRNVVFDIFTGKGQDMVFT    120
OsC4H1/2a/2b-TKO-b         LARRFGDILLLRMGVRNLVVVSSPDLAKEVLHTQGVEFGSRTRNVVFDIFTGKGQDMVFT    120

WT                         VYGDHWRKMRRIMTVPFFTNKVVAQNRAGWEEEARLVVEDVRRDPAAATSGVVIRRRLQL    180
OsC4H1-KO                  VYGDHWRKMRRIMTVPFFTNKVVAQNRAGWEEEARLVVEDVRRDPAAATSGVVIRRRLQL    180
OsC4H1/2a/2b-TKO-a         VYGDHWRKMRRIMTVPFFTNKVVAQNRAGWEEEARLVVEDVRRDPAAATSGVVIRRRLQL    180
OsC4H1/2a/2b-TKO-b         VYGDHWRKMRRIMTVPFFTNKVVAQNRAGWEEEARLVVEDVRRDPAAATSGVVIRRRLQL    180

WT                         MMYNDMFRIMFDRRFDSVDDPLFNKLKAFNAERSRLSQSFEYNYGDFIPVLRPFLRRYLA    240
OsC4H1-KO                  MMYNDMFRIMFDRRFDSVDDPLFNKLKAFNAERSRLSQSFEYNYGDFIPVLRPFLRRYLA    240
OsC4H1/2a/2b-TKO-a         MMYNDMFRIMFDRRFDSVDDPLFNKLKAFNAERSRLSQSFEYNYGDFIPVLRPFLRRYLA    240
OsC4H1/2a/2b-TKO-b         MMYNDMFRIMFDRRFDSVDDPLFNKLKAFNAERSRLSQSFEYNYGDFIPVLRPFLRRYLA    240

WT                         RCHQLKSQRMKLFEDHFVQERKRVMEQTGEIRCAMDHILEAERKGEINHDNVLYIVENIN    300
OsC4H1-KO                  RCHQLKSQRMKLFEDHFVQERKRVMEQTGEIRCAMDHILEAERKGEINHDNVLYIVENIN    300
OsC4H1/2a/2b-TKO-a         RCHQLKSQRMKLFEDHFVQERKRVMEQTGEIRCAMDHILEAERKGEINHDNVLYIVENIN    300
OsC4H1/2a/2b-TKO-b         RCHQLKSQRMKLFEDHFVQERKRVMEQTGEIRCAMDHILEAERKGEINHDNVLYIVENIN    300
                      oxygen binding region
WT                         VAAIETTLWSIEWGIAELVNHPSIQSKVREEMASVLGGAAVTEPDLERLPYLQAVVKETL    360
OsC4H1-KO                  VAAIETTLWSIEWGIAELVNHPSIQSKVREEMASVLGGAAVTEPDLERLPYLQAVVKETL    360
OsC4H1/2a/2b-TKO-a         VAAIETTLWSIEWGIAELVNHPSIQSKVREEMASVLGGAAVTEPDLERLPYLQAVVKETL    360
OsC4H1/2a/2b-TKO-b         VAAIETTLWSIEWGIAELVNHPSIQSKVREEMASVLGGAAVTEPDLERLPYLQAVVKETL    360

WT                         RLRMAIPLLVPHMNLADGKLAGYDIPAESKILVNAWFLANDPKRWVRPDEFRPERFLEEE    420
OsC4H1-KO                  RLRMAIPLLVPHMNLADGKLAGYDIPAESKILVNRVVPRQRPQAVGAPRRV*AGEVPG--    417
OsC4H1/2a/2b-TKO-a         RLRMAIPLLVPHMNLADGKLAGYDIPAESKILVNRVVPRQRPQAVGAPRRV*AGEVPG--    417
OsC4H1/2a/2b-TKO-b         RLRMAIPLLVPHMNLADGKLAGYDIPAESKILVNSVVPRQRPQAVGAPRRV*AGEVPG--    417
                                heme binding region
WT                         KAVEAHGNDFRFVPFGVGRRSCPGIILALPIIGITLGRLVQSFDLLPPPGMDKV-DTTEK    479
OsC4H1-KO                  -GGEGRGGARQRLPLRALRRRPPQLPRDHPR-------AAHHRDHARPPRPELRPAAAAR    469
OsC4H1/2a/2b-TKO-a         -GGEGRGGARQRLPLRALRRRPPQLPRDHPR-------AAHHRDHARPPRPELRPAAAAR    469
OsC4H1/2a/2b-TKO-b         -GGEGRGGARQRLPLRALRRRPPQLPRDHPR-------AAHHRDHARPPRPELRPAAAAR    469

WT                         PGQFSNQILKHATVVCKPIDA*---------------         500
OsC4H1-KO                  DGQGGH------HREARPVQQPDPQARHRRLQAHRRL         500
OsC4H1/2a/2b-TKO-a         DGQGGH------HREARPVQQPDPQARHRRLQAHRRL         500
OsC4H1/2a/2b-TKO-b         DGQGGH------HREARPVQQPDPQARHRRLQAHRRL         500
```

**Supplemental Figure S2. Predicted effects of CRISPR/Cas9-induced mutations on OsC4H1.**

Conserved cytochrome P450 motifs are highlighted. Red asterisks indicate the first premature stop codons after the mutation site. WT, wild type control line; *OsC4H1*-KO, *OsC4H1* single-knockout line; *OsC4H1/2a/2b*-TKO-a and *OsC4H1/2a/2b*-TKO-b, *OsC4H1*, *OsC4H2a* and *OsC4H2b* triple-knockout lines.

>OsC4H2a

```
WT                      MAASVVRVAIATGASLAVHLFVKSFLQAQHPALTLLLPVAVFAGIAVGAKGGNGGDGKAP   60
OsC4H2a/2b-DKO          MAASVVRVAIATGASLAVHLFVKSFLQAQHPALTLLLPVAVFAGIAGGREGRERGRREGA   60
OsC4H1/2a/2b-TKO-a      MAASVVRVAIATGASLAVHLFVKSFLQAQHPALTLLLPVAVFAGIAGGREGRERGRREGA   60
OsC4H1/2a/2b-TKO-b      MAASVVRVAIATGASLAVHLFVKSFLQAQHPALTLLLPVAVFAGIAGGREGRERGRREGA   60
                      cluster of proline
WT                      PGPAAVPVFGNWLHVGNDLNHRFLAAMSARYGPVFRLRLGVRNLVVVSDPKLAT------   114
OsC4H2a/2b-DKO          AGAGGRAGV------------RQLAARRERPEPQ--VPRGDVGAVRARVPSAAGRAQPGG   106
OsC4H1/2a/2b-TKO-a      AGAGGRAGV------------RQLAARRERPEPQ--VPRGDVGAVRARVPSAAGRAQPGG   106
OsC4H1/2a/2b-TKO-b      AGAGGRAGV------------RQLAARRERPEPQ--VPRGDVGAVRARVPSAAGRAQPGG   106

WT                      ---------------------------EVLHTQGVEFG----SRPRNVV---FDIFTANG   140
OsC4H2a/2b-DKO          GVGPEAGDGGAAHAGRGVRLPPAQRRLRHLHRQRRRHGVHRVRRPLATHAPRHDAAVLHG   166
OsC4H1/2a/2b-TKO-a      GVGPEAGDGGAAHAGRGVRLPPAQRRLRHLHRQRRRHGVHRVRRPLATHAPRHDAAVLHG   166
OsC4H1/2a/2b-TKO-b      GVGPEAGDGGAAHAGRGVRLPPAQRRLRHLHRQRRRHGVHRVRRPLATHAPRHDAAVLHG   166

WT                      AD--MVFTEYGD--HWRRMRRVMTLPFFTARVVQQYKAMWEAEMDAVVDDVRGDAVAQGT   196
OsC4H2a/2b-DKO          ARRAAVQGHVGGRDGRRRGRRARRRGGAGHRLRGATQAAAHAVQHHVPDDVRRAVRVGGR   226
OsC4H1/2a/2b-TKO-a      ARRAAVQGHVGGRDGRRRGRRARRRGGAGHRLRGATQAAAHAVQHHVPDDVRRAVRVGGR   226
OsC4H1/2a/2b-TKO-b      ARRAAVQGHVGGRDGRRRGRRARRRGGAGHRLRGATQAAAHAVQHHVPDDVRRAVRVGGR   226

WT                      GFVVRRRLQLMLYNIMYRMMFDARFESVDDPMFIEATRFNSERSRLAQSFEYNYGDFIPI   256
OsC4H2a/2b-DKO          PHV-HRGHQVQLRAQPPRAE-----------------------------LRVQLRRLHPH   256
OsC4H1/2a/2b-TKO-a      PHV-HRGHQVQLRAQPPRAE-----------------------------LRVQLRRLHPH   256
OsC4H1/2a/2b-TKO-b      PHV-HRGHQVQLRAQPPRAE-----------------------------LRVQLRRLHPH   256

WT                      LRPFLRGYLNKCRDLQSRRLAFFNNNYVEKR------------RKVMDTP----GDRNKL   300
OsC4H2a/2b-DKO          PPSLLAGLPQQVP*PPEQEARLLQQQLRREEKEGDGHSGRQEQAPVRDRPYP*GGEERRA   314
OsC4H1/2a/2b-TKO-a      PPSLLAGLPQQVP*PPEQEARLLQQQLRREEKEGDGHSGRQEQAPVRDRPYP*GGEERRA   314
OsC4H1/2a/2b-TKO-b      PPSLLAGLPQQVP*PPEQEARLLQQQLRREEKEGDGHSGRQEQAPVRDRPYP*GGEERRA   314
                                                        oxygen binding region
WT                      RCAIDHILEAE-KNGELTAENVIYIVENIN----VAAIET-----TLWSIEWALAEVVNH   350
OsC4H2a/2b-DKO          DGGERDLHRGEHQRGRHRDDALVHRVGAGRGRQPPGGAEQGPRRDQRRARRRRAHHRVQH   374
OsC4H1/2a/2b-TKO-a      DGGERDLHRGEHQRGRHRDDALVHRVGAGRGRQPPGGAEQGPRRDQRRARRRRAHHRVQH   374
OsC4H1/2a/2b-TKO-b      DGGERDLHRGEHQRGRHRDDALVHRVGAGRGRQPPGGAEQGPRRDQRRARRRRAHHRVQH   374

WT                      PAVQSKVRAEINDVLG-DDEPITESSIHKLTYLQAVIKETLRLHSPIPL-----------   398
OsC4H2a/2b-DKO          PQADLPAGRDQGDAAAALPDPAAG-AAHEP-----GGGQARRVHHPQGIQGGGERVVAGQ   428
OsC4H1/2a/2b-TKO-a      PQADLPAGRDQGDAAAALPDPAAG-AAHEP-----GGGQARRVHHPQGIQGGGERVVAGQ   428
OsC4H1/2a/2b-TKO-b      PQADLPAGRDQGDAAAALPDPAAG-AAHEP-----GGGQARRVHHPQGIQGGGERVVAGQ   428

WT                      ---------LVPHMN-L--------EEAKLGGYTIPKGSKVVVNAWWLANNPALWENPEE   440
OsC4H2a/2b-DKO          QPGAVGEPRGVPA*AVLGEGERRGRHRRREGGLQVP----------------ALRRGPPQ   471
OsC4H1/2a/2b-TKO-a      QPGAVGEPRGVPA*AVLGEGERRGRHRRREGGLQVP----------------ALRRGPPQ   471
OsC4H1/2a/2b-TKO-b      QPGAVGEPRGVPA*AVLGEGERRGRHRRREGGLQVP----------------ALRRGPPQ   471
                                                                heme binding region
WT                      FRPERFL-------EKESGV---DATVAGK-----VDFR-F--LP--FGVGRRSCPGIIL   480
OsC4H2a/2b-DKO          LPGDHPGAAHPGAHRREAGEELRDGAAAGRGEAGRERERRAVQPPHRQALRRRLPPHLCL   531
OsC4H1/2a/2b-TKO-a      LPGDHPGAAHPGAHRREAGEELRDGAAAGRGEAGRERERRAVQPPHRQALRRRLPPHLCL   531
OsC4H1/2a/2b-TKO-b      LPGDHPGAAHPGAHRREAGEELRDGAAAGRGEAGRERERRAVQPPHRQALRRRLPPHLCL   531

WT                      ALPILALIVGKLVRSFEMVPPPGVEKLDVSEKGGQFSLHIAKHSVVAFHPISA*      533
OsC4H2a/2b-DKO          ------------------------------------------------------      531
OsC4H1/2a/2b-TKO-a      ------------------------------------------------------      531
OsC4H1/2a/2b-TKO-b      ------------------------------------------------------      531
```

**Supplemental Figure S3. Predicted effects of CRISPR/Cas9-induced mutations on OsC4H2a.**

Conserved cytochrome P450 motifs are highlighted. Red asterisks indicate the first premature stop codons after the mutation site. WT, wild type control line; *OsC4H2a/2b*-DKO, *OsC4H2a* and *OsC4H2b* double-knockout line; *OsC4H1/2a/2b*-TKO-a and *OsC4H1/2a/2b*-TKO-b, *OsC4H1*, *OsC4H2a* and *OsC4H2b* triple-knockout lines.

>OsC4H2b

```
WT                        MAASAMRVAIATGASLAVHLFVKSFVQAQHPALTLLLPVAVFVGIAVGAKGGSGGDGK--   58
OsC4H2a/2b-DKO            MAASAMRVAIATGASLAVHLFVKSFVQAQHPALTLLLPVAVFVGIAGGREGRERW*REGA   59
OsC4H1/2a/2b-TKO-a        MAASAMRVAIATGASLAVHLFVKSFVQAQHPALTLLLPVAVFVGIAGGREGRERW*REGA   59
OsC4H1/2a/2b-TKO-b        MAASAMRVAIATGASLAVHLFVKSFVQAQHPALTLLLPVAVFVGIAGGREGRERW*REGA   59
                                                   cluster of proline
WT                        -------------------APPGPAA--------VPVFGNWLQVGNDLNHRFLAAMSARY   91
OsC4H2a/2b-DKO            AGAGGRAGVRQLAAGGQRPEPPVPRGDVGTVRSRVPSAAGRAQPGGGVGPEAGDGGAAHA   119
OsC4H1/2a/2b-TKO-a        AGAGGRAGVRQLAAGGQRPEPPVPRGDVGTVRSRVPSAAGRAQPGGGVGPEAGDGGAAHA   119
OsC4H1/2a/2b-TKO-b        AGAGGRAGVRQLAAGGQRPEPPVPRGDVGTVRSRVPSAAGRAQPGGGVGPEAGDGGAAHA   119

WT                        GPVFRLRLGVRNL------------------VVVSDPKLATEVLHTQGVEFGSRPRNVVF   133
OsC4H2a/2b-DKO            GRGVRLPPAQRRLRHLHRQRRRHGVHRVRRPLATHAPRHDAAVLH------GARRAAVQG   173
OsC4H1/2a/2b-TKO-a        GRGVRLPPAQRRLRHLHRQRRRHGVHRVRRPLATHAPRHDAAVLH------GARRAAVQG   173
OsC4H1/2a/2b-TKO-b        GRGVRLPPAQRRLRHLHRQRRRHGVHRVRRPLATHAPRHDAAVLH------GARRAAVQG   173

WT                        DIFTANGADMVFTEYGDHWRRMRRVMTLPFFTARVVQQYKAMWEAEMDAVVDDVRGDAVA   193
OsC4H2a/2b-DKO            HVG---GR--------DGRRRGRRARRRGGAGHRLRGATQAAAHAVQHHVPDDVRRAVRV   222
OsC4H1/2a/2b-TKO-a        HVG---GR--------DGRRRGRRARRRGGAGHRLRGATQAAAHAVQHHVPDDVRRAVRV   222
OsC4H1/2a/2b-TKO-b        HVG---GR--------DGRRRGRRARRRGGAGHRLRGATQAAAHAVQHHVPDDVRRAVRV   222

WT                        QGTGFVVRRRLQLMLYNIMYRMMFDARFESVDDPMFIEATRFNSERSRLAQSFEYNYGDF   253
OsC4H2a/2b-DKO            GGRPHV-HRGHQVQLRAQPPRAE-----------------------------LRVQLRRL   252
OsC4H1/2a/2b-TKO-a        GGRPHV-HRGHQVQLRAQPPRAE-----------------------------LRVQLRRL   252
OsC4H1/2a/2b-TKO-b        GGRPHV-HRGHQVQLRAQPPRAE-----------------------------LRVQLRRL   252

WT                        IPILRPFLRGYLNKCRDLQSRRLAFFNNNYVEKR------------RKVMDTP----GDR   297
OsC4H2a/2b-DKO            HPHPPSLLAGLPQQVP*PPEQEARLLQQQLRREEKEGDGHSGRQEQAPVRDRPYP*GGEE   310
OsC4H1/2a/2b-TKO-a        HPHPPSLLAGLPQQVP*PPEQEARLLQQQLRREEKEGDGHSGRQEQAPVRDRPYP*GGEE   310
OsC4H1/2a/2b-TKO-b        HPHPPSLLAGLPQQVP*PPEQEARLLQQQLRREEKEGDGHSGRQEQAPVRDRPYP*GGEE   310
                                                                oxygen binding region
WT                        NKLRCAIDHILEAE-KNGELTAENVIYIVENIN----VAAIET-----TLWSIEWALAEV   347
OsC4H2a/2b-DKO            RRADGGERDLHRGEHQRGRHRDDALVHRVGAGRGRQPPGGAEQGPRRDQRRARRRRAHHR   370
OsC4H1/2a/2b-TKO-a        RRADGGERDLHRGEHQRGRHRDDALVHRVGAGRGRQPPGGAEQGPRRDQRRARRRRAHHR   370
OsC4H1/2a/2b-TKO-b        RRADGGERDLHRGEHQRGRHRDDALVHRVGAGRGRQPPGGAEQGPRRDQRRARRRRAHHR   370

WT                        VNHPAVQSKVRAEINDVLG-DDEPITESSIHKLTYLQAVIKETLRLHSPIPL--------   398
OsC4H2a/2b-DKO            VQHPQADLPAGRDQGDAAAALPDPAAG-AAHEP-----GGGQARRVHHPQGIQGGGERVV   424
OsC4H1/2a/2b-TKO-a        VQHPQADLPAGRDQGDAAAALPDPAAG-AAHEP-----GGGQARRVHHPQGIQGGGERVV   424
OsC4H1/2a/2b-TKO-b        VQHPQADLPAGRDQGDAAAALPDPAAG-AAHEP-----GGGQARRVHHPQGIQGGGERVV   424

WT                        ------------LVPHMN-L--------EEAKLGGYTIPKGSKVVVNAWWLANNPALWEN   437
OsC4H2a/2b-DKO            AGQQPGAVGEPRGVPA*AVLGEGERRGRHRRREGGLQVP----------------ALRRG   467
OsC4H1/2a/2b-TKO-a        AGQQPGAVGEPRGVPA*AVLGEGERRGRHRRREGGLQVP----------------ALRRG   467
OsC4H1/2a/2b-TKO-b        AGQQPGAVGEPRGVPA*AVLGEGERRGRHRRREGGLQVP----------------ALRRG   467

                                                                         heme binding region
WT                        PEEFRPERFL-------EKESGV---DATVAGK-----VDFR-F--LP--FGVGRRSCPG   477
OsC4H2a/2b-DKO            PPQLPGDHPGAAHPGAHRREAGEELRDGAAAGRGEAGRERERRAVQPPHRQALRRRLPPH   527
OsC4H1/2a/2b-TKO-a        PPQLPGDHPGAAHPGAHRREAGEELRDGAAAGRGEAGRERERRAVQPPHRQALRRRLPPH   527
OsC4H1/2a/2b-TKO-b        PPQLPGDHPGAAHPGAHRREAGEELRDGAAAGRGEAGRERERRAVQPPHRQALRRRLPPH   527

WT                        IILALPILALIVGKLVRSFEMVPPPGVEKLDVSEKGGQFSLHIAKHSVVAFHPISA*   533
OsC4H2a/2b-DKO            LCL------------------------------------------------------   530
OsC4H1/2a/2b-TKO-a        LCL------------------------------------------------------   530
OsC4H1/2a/2b-TKO-b        LCL------------------------------------------------------   530
```

**Supplemental Figure S4. Predicted effects of CRISPR/Cas9-induced mutations on OsC4H2b.**

Conserved cytochrome P450 motifs are highlighted. Red asterisks indicate the first premature stop codons after the mutation site. WT, wild type control line; *OsC4H2a/2b*-DKO, *OsC4H2a* and *OsC4H2b* double-knockout line; *OsC4H1/2a/2b*-TKO-a and *OsC4H1/2a/2b*-TKO-b, *OsC4H1*, *OsC4H2a* and *OsC4H2b* triple-knockout lines.

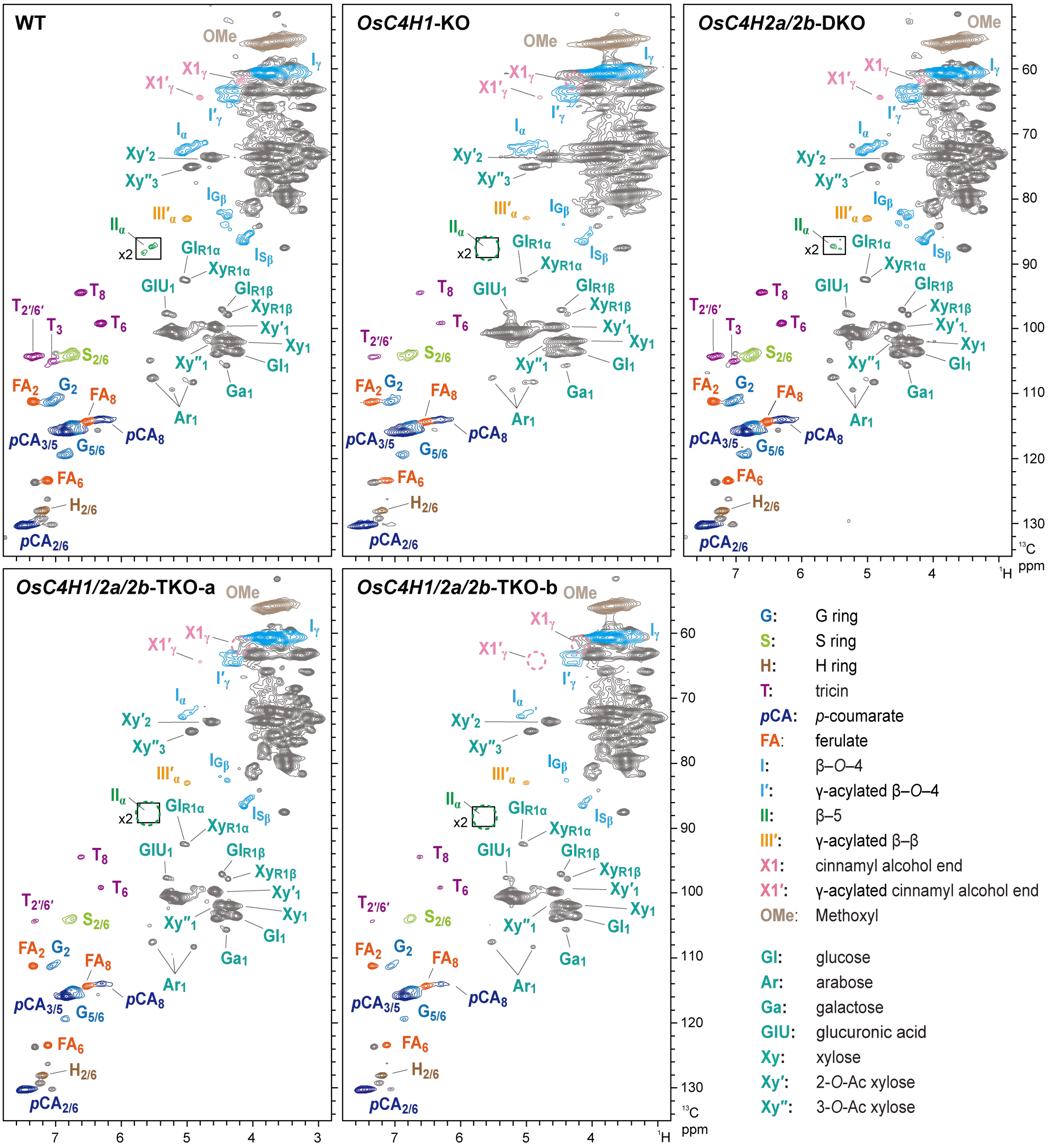


**Supplemental Figure S5. 2D HSQC NMR spectra of *C4H*-knockout rice culm cell walls.**

The $^{1}H$–$^{13}C$ short-range correlation (HSQC) NMR spectra of ball-milled whole rice culm cell walls. Signal assignments are listed in **Supplemental Table S6**. Boxes labelled ×2 represent regions with the scale vertically enlarged by 2-fold. NMR analysis was conducted for culm cell wall residue (CWR) samples pooled from three biologically independent plants for each line. WT, wild type control line; *OsC4H1*-KO, *OsC4H1* single-knockout line; *OsC4H2a/2b*-DKO, *OsC4H2a* and *OsC4H2b* double-knockout line; *OsC4H1/2a/2b*-TKO-a and *OsC4H1/2a/2b*-TKO-b, *OsC4H1*, *OsC4H2a* and *OsC4H2b* triple-knockout lines.

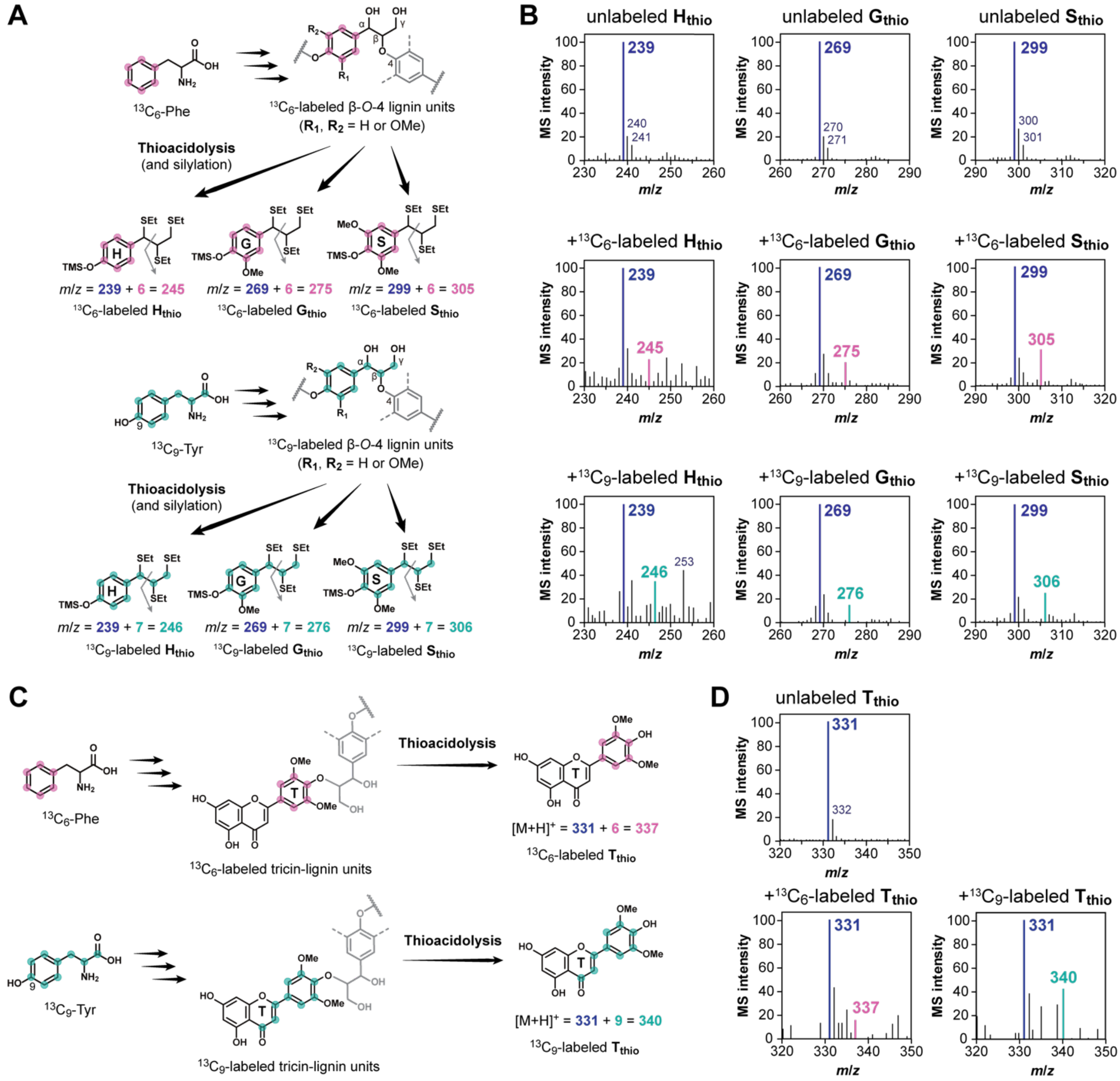

**Supplemental Figure S6. Analysis of $^{13}$C-labeled lignin units by thioacidolysis.**

**A** and **C**) Schemes of $^{13}$C-labeled Phe and Tyr incorporation into monolignol-derived (**A**) and tricin-derived (**C**) lignin units, and their release by thioacidolysis. **B** and **D**) MS spectra of the thioacidolysis-derived monomeric products released from monolignol-derived ($\mathbf{H_{thio}}$, $\mathbf{G_{thio}}$, and **Sthio** detected by GC-MS after silylation) (**B**) and tricin-derived ($\mathbf{T_{thio}}$ detected by LC-MS) (**D**) lignin units. Major ion peaks derived from unlabeled and $^{13}$C-labeled products are highlighted.

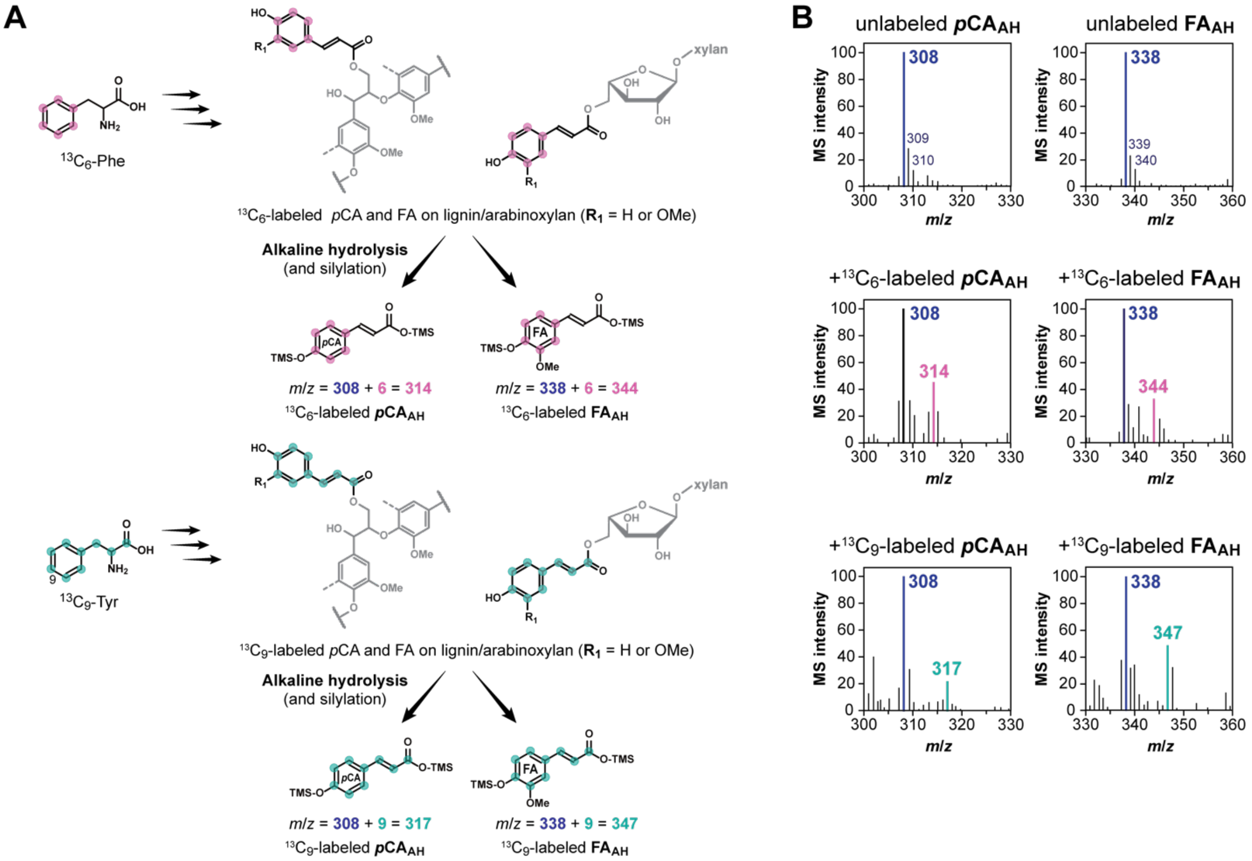


**Supplemental Figure S7. Analysis of $^{13}$C-labeled cell wall-bound hydroxycinnamates by alkaline hydrolysis.**

**A)** Schemes of $^{13}$C-labeled Phe and Tyr incorporation into lignin- and arabinoxylan-bound p-coumarate (*p*CA) and ferulate (FA), and their release by mild alkaline hydrolysis. **B**) MS spectra of *p*CA and FA released from cell wall-bound *p*CA and FA (***p*CA$_{AH}$** and **FA$_{AH}$** detected by GC-MS after silylation). Major ion peaks derived from unlabeled and 13C-labeled pCAAH and FAAH are highlighted.

**Supplemental Table S1. Gene locus and accession numbers of C4H homologs examined.**

| Species | C4H isoform names | Gene locus | Protein accession |
|---|---|---|---|
| *O. sativa* | OsC4H1 (CYP73A38) | LOC_Os05g25640 | XP_015639656 |
| | CYP73A35p | LOC_Os01g60450 | XP_015635394 |
| | OsC4H2a (CYP73A39) | LOC_Os02g26770 | XP_015623447 |
| | OsC4H2b (CYP73A40) | LOC_Os02g26810 | XP_015626579 |
| *B. distachyon* | BdC4H1a (CYP73A92) | Bradi2g31510 | XP_003568699 |
| | BdC4H1b (CYP73A93) | Bradi2g53470 | XP_003564495 |
| | BdC4H2 (CYP73A94) | Bradi3g43160 | XP_003574953 |
| *Z. mays* | ZmC4H1a | GRMZM2G139874 | NP_001140726 |
| | ZmC4H1b | GRMZM2G147245 | NP_001140681 |
| | ZmC4H1c | GRMZM2G010468 | XP_008657127 |
| | ZmC4H2 | GRMZM2G028677 | NP_001151365 |
| *S. bicolor* | SbC4H1a | Sobic.002G126600 | XP_002461939 |
| | SbC4H1b | Sobic.003G337400 | XP_002458683 |
| | SbC4H2 | Sobic.004G141200 | XP_002452044 |
| *N. tabacum* | NtC4H1a | Nitab4.5_0005901g0010 | XP_016493546 |
| | NtC4H1b | Nitab4.5_0002902g0020 | NP_001313097 |
| | NtC4H1c | Nitab4.5_0000105g0250 | NP_001312445 |
| | NtC4H2a | Nitab4.5_0005105g0020 | NP_001312254 |
| | NtC4H2b | Nitab4.5_0000404g0200 | NP_001313000 |
| *A. thaliana* | AtC4H1/REF3 (CYP73A5) | AT2G30490 | NP_180607 |
| *P. trichocarpa* | PtrC4H1a | Potri.013G157900 | XP_002319974 |
| | PtrC4H1b | Potri.019G130700 | XP_002319974 |
| | PtrC4H2a | Potri.018G146100 | XP_024446000 |
| *M. truncatula* | MtrC4H1a | Medtr5g075450 | XP_003616037 |
| | MtrC4H2a | Medtr1g111240 | XP_013470209 |

**Supplemental Table S2. Amino acid sequence identity of rice and Arabidopsis C4H members.**

| C4H protein | Class I C4H | | | Class II C4H | |
|---|---|---|---|---|---|
| | AtC4H1/REF3 (CYP73A5) | CYP73A35p | OsC4H1 (CYP73A38) | OsC4H2a (CYP73A39) | OsC4H2b (CYP73A40) |
| AtC4H1/REF3 (CYP73A5) | - | 74% | 76% | 65% | 62% |
| CYP73A35p | 74% | - | 69% | 58% | 60% |
| OsC4H1 (CYP73A38) | 76% | 69% | - | 62% | 64% |
| OsC4H2a (CYP73A39) | 65% | 58% | 62% | - | 99% |
| OsC4H2b (CYP73A40) | 62% | 60% | 60% | 99% | - |

**Supplemental Table S3. Oligonucleotide sequences used for constructing sgRNAs.**

| Target genes | sgRNA# | Exon # | Target +PAM | GC content | CRISPR-P score* | Off-target score* |
|---|---|---|---|---|---|---|
| *OsC4H1* | 1 | 3 | TCTCCGGCCTAAACTCGTCG**GGG** | 60% | 0.7 | 0.161 |
| | 2 | 3 | CCAAGATCCTGGTGAACGCG**TGG** | 60% | 0.5 | 0.284 |
| *OsC4H2a/2b* | 1 | 1 | CCAGTTGCCGAACACCGGCA**CGG** | 65% | 0.7 | 0.59 |
| | 2 | 1 | GTGTTTGTCGGCATTGCGGT**GGG** | 55% | 0.5 | 0.368 |

*Determined by CRISPR-P 2.0 (Liu et al. 2017).

### Supplemental Table S4. Off-target analyses of *C4H*-knockout rice mutants.

| sgRNA target | Off-target site name | Sequence (5′ to 3′)[a] | Off-target score[b] | Location (Chromosome: start) | Region | Mutation[c] |
|---|---|---|---|---|---|---|
| *OsC4H1* guide 2 (target 1) | site 1-1 | TCTCCGGCATGAACTTGTCAGGG | 0.161 | 3:+7832377 | CDS (LOC_Os03g14400) | n.d. |
| | site 1-2 | TCTCCCGCCAAAACTCTTCAAGG | 0.137 | 5:+22238695 | intergenic | n.d. |
| | site 1-3 | TCTCCGGCCGGAGCTCGCCGCGG | 0.091 | 1:-1570640 | CDS (LOC_Os01g03750) | n.d. |
| OsC4H1 guide 6 (target 2) | site 2-1 | CCAAGGTGGTGGTGAACGCGTGG | 0.284 | 2:-15717380 | CDS (LOC_Os02g26770) | n.d. |
| | site 2-2 | CCAAGGTGGTGGTGAACGCGTGG | 0.284 | 2:+15736323 | CDS (LOC_Os02g26810) | n.d. |
| | site 2-3 | GCAAGAGGCTGGTGAACTCGAGG | 0.230 | 11:-28934483 | CDS (LOC_Os11g47970) | n.d. |
| *OsC4H2a/2b* guide 2 (target 1) | site 3-1 | CGAGTCACCGAACACCAGCATGG | 0.590 | 2:+547603 | CDS (LOC_Os02g01980) | n.d. |
| | site 3-2 | CCAGTCACCGAACTCCGGCGAGG | 0.371 | 3:-4193243 | CDS (LOC_Os03g08220) | n.d. |
| | site 3-3 | CCAGTCACCGAACTCCGGCGAGG | 0.321 | 10:+2455530 | CDS (LOC_Os10g05020) | n.d. |
| *OsC4H2a/2b* guide 6 (target 1) | site 4-1 | GGGTTTACTGGCATTGCTGTTGG | 0.490 | 9:+19754854 | CDS (LOC_Os09g33500) | n.d. |
| | site 4-2 | GTGTCAGGCGGCATAGCGGTGGG | 0.323 | 11:-20264554 | CDS (LOC_Os11g34624) | n.d. |
| | site 4-3 | GTGTTTGTCGGCGTTGTGGTGGG | 0.266 | 12:-20899361 | CDS (LOC_Os12g34524) | n.d. |

[a]Underlined: mismatch nucleotides compared with the sequence of sgRNA. [b]Determined by CRISPR-P 2.0 (Liu et al. 2017). CDS: coding sequence. n.d.: not detected.

**Supplemental Table S5. Neutral sugar analysis of *C4H*-knockout rice culm cell walls.**

| Component (mg/g CWR) | WT | *OsC4H1-KO* | *OsC4H2a/2b*-DKO | *OsC4H1/2a/2b-TKO-a* | *OsC4H1/2a/2b*-TKO-b |
|---|---|---|---|---|---|
| Crystalline glucose[1] | 433.57 ± 25.7 [a] | 407.2 ± 10.7 [a] | 435.4 ± 13.6 [a] | 431.6 ± 18.1 [a] | 412.8 ± 21.6 [a] |
| Amorphous glucose[2] | 33.3 ± 6.5 [b] | **47.8 ± 3.9 [a]** | **50.7 ± 5.1 [a]** | **49.6 ± 5.1 [a]** | 46.9 ± 4.8 [ab] |
| Xylose | 71.5 ± 9.1 [a] | 77.9 ± 5.4 [a] | 81.5 ± 7.4 [a] | 82.4 ± 1.8 [a] | 78.0 ± 4.0 [a] |
| Arabinose | 22.3 ± 3.0 [a] | 25.1 ± 1.5 [a] | 25.1 ± 2.6 [a] | 26.1 ± 1.6 [a] | 24.6 ± 1.1 [a] |
| Mannose | 4.4 ± 0.4 [a] | 5.1 ± 0.5 [a] | 4.9 ± 0.5 [a] | 4.9 ± 0.5 [a] | 4.3 ± 0.5 [a] |
| Galactose | 16.8 ± 2.9 [a] | 18.8 ± 0.7 [a] | 17.9 ± 2.2 [a] | 18.4 ± 2.3 [a] | 17.4 ± 1.2 [a] |

[1]Glucose released from the trifluoroacetic acid-insoluble cell wall fractions. [2]Glucose released from the trifluoroacetic acid-soluble cell wall fractions. Values are mean ± standard deviation ($n = 6$ for 5-month-old mature plants; $n = 15$ for 4-week-old plantlets). Different letters on values indicate significant differences and significant differences over wild-type control are highlighted in bold (one-way ANOVA with Tukey's test, $P < 0.05$). CWR, cell wall residue. WT, wild type control line; *OsC4H1*-KO, *OsC4H1* single-knockout line; *OsC4H2a/2b*-DKO, *OsC4H2a* and *OsC4H2b* double-knockout line; *OsC4H1/2a/2b*-TKO-a and *OsC4H1/2a/2b*-TKO-b, *OsC4H1*, *OsC4H2a* and *OsC4H2b* triple-knockout lines.

**Supplemental Table S6. Peak assignments in 2D HSQC NMR spectra of rice cell walls.**

| Labels | $\delta_C/\delta_H$ (ppm) | Assignment |
|---|---|---|
| *Lignin and hydroxycinnamate signals* | | |
| **$S_{2/6}$** | 103.6/6.76 | C2–H2 and C6–H6 in syringyl units |
| **$G_2$** | 103.9/6.81 | C2–H2 in guaiacyl units |
| **$G_{5/6}$** | 119..2/6.86 | C5–H5 and C6–H6 in guaiacyl units |
| **$H_{2/6}$** | 127.9/7.21 | C2–H2 and C6–H6 in *p*-hydroxyphenyl units |
| **$T_3$** | 105.1/7.08 | C3–H3 in tricin residues |
| **$T_6$** | 99.9/6.22 | C6–H6 in tricin residues |
| **$T_8$** | 94.8/6.53 | C8–H8 in tricin residues |
| **$T_{2'/6'}$** | 104.0/7.32 | C2′–H2′ and C6′–H6′ in tricin residues |
| ***p*$CA_{2/6}$** | 130.0/7.44 | C2–H2 and C6–H6 in *p*-coumarate residues |
| ***p*$CA_{3/5}$** | 111.2/7.35 | C3–H3 and C5–H5 in *p*-coumarate residues |
| ***p*$CA_8$** | 113.8/6.26 | C8–H8 in *p*-coumarate residues |
| **$FA_2$** | 112.6/7.36 | C2–H2 in ferulate residues |
| **$FA_6$** | 123.1/7.14 | C6–H6 in ferulate residues |
| **$FA_8$** | 114.1/6.54 | C8–H8 in ferulate residues |
| **$I_\alpha$** | 72.1/4.97 | Cα–Hα in β–*O*–4 units |
| **$I_\beta$** | 86.2/4.16, 82.7/4.40 | Cβ–Hβ in β–*O*–4 units |
| **$I_\gamma$** | 60.4/3.71 | Cγ–Hγ in γ-free β–*O*–4 units |
| | 64.2/4.43 | Cγ–Hγ in γ-acylated β–*O*–4 units |
| **$II_\alpha$** | 87.4/5.55 | Cα–Hα in β–5 substructures |
| **$II_\beta$** | 53.2/3.52 | Cβ–Hβ in β–5 substructures |
| **$III_\alpha$** | 83.1/5.01 | Cα–Hα in tetrahydrofuran-type β–β substructures |
| **$X1_\gamma$** | 61.9/4.15 | Cγ–Hγ in γ-free cinnamyl alcohol end-units |
| **$X1'_\gamma$** | 64.4/4.82 | Cγ–H in γ-acylated cinnamyl alcohol end-units |
| **OMe** | 55.9/3.70 | C–H in aromatic methoxyl groups |
| *Polysaccharide signals* | | |
| **$Gl_1$** | 103.5/4.24 | C1–H1 in (1→4)-β-D-glucopyranosyl units |
| **$X_1$** | 101.7/4.26 | C1–H1 in (1→4)-β-D-xylopyranosyl units |
| **$X'_1$** | 99.7/4.50 | C1–H1 in 2-*O*-acetyl-β-D-xylopyranosyl units |
| **$X'_2$** | 73.6/4.67 | C2–H2 in 2-*O*-acetyl-β-D-xylopyranosyl units |
| **$X''_1$** | 101.8/4.87 | C1–H1 in 3-*O*-acetyl-β-D-xylopyranosyl units |
| **$X''_3$** | 75.2/4.97 | C3–H3 in 3-*O*-acetyl-β-D-xylopyranosyl units |
| **$A_1$** | 109.3/5.23, 108.1/4.92, 107.7/5.06 | C1–H1 in α-L-arabinofuranosyl units |
| **$Ga_1$** | 105.5/4.41 | C1–H1 in (1→4)-β-D-galactopyranosyl units |
| **$GlU_1$** | 97.5/5.31 | C1–H1 in 4-*O*-methyl-α-D-glucuronopyranosyl units |

Measured in DMSO-$d_6$/Py-$d_5$ (4:1, v/v). Assignment was based on comparison with literature data reported in Kim and Ralph (2010), Mansfield et al. (2012), Kim et al. (2017), Tobimatsu et al. (2019), Afifi et al. (2022), and Martin et al. (2023).

**Supplemental Table S7. Primers and oligonucleotides used in this study.**

| Purpose | Sequence |
|---|---|
| RTq-PCR of CYP73A35p | 5′-CGCAGAGCTTCGAGTACAAC-3′<br>5′-TTCATGCGCTGGGACTTGAG-3′ |
| RTq-PCR of OsC4H1 | 5′-TCATGACGGTGCCGTTCTTC-3′<br>5′-AGGATGTGATCAATGGCGCA-3′ |
| RTq-PCR of OsC4H2a and OsC4H2b | 5′-CATCCTGCTCTCACCTTGCT-3′<br>5′-TGCAGCCAGTTGCCGAA-3′ |
| Generation of *OsC4H1-KO* and *OsC4H1/2a/2b-TKO* (target site 1) | 5′-GTTGCCAAGATCCTGGTGAACGCG-3′<br>5′-AAACCGCGTTCACCAGGATCTTGG-3′ |
| Generation of *OsC4H1-KO* and *OsC4H1/2a/2b-TKO* (target site 2) | 5′-GTTGTCTCCGGCCTAAACTCGTCG-3′<br>5′-AAACCGACGAGTTTAGGCCGGAGA-3′ |
| Generation of *OsC4H2a/2b-DKO KO* and *OsC4H1/2a/2/b-TKO* (target site 1) | 5′-GTTGGTGTTTGTCGGCATTGCGGT-3′<br>5′-AAACACCGCAATGCCGACAAACAC-3′ |
| Generation of *OsC4H2a/2b-DKO KO* and *OsC4H1/2a/2/b-TKO* (target site 2) | 5′-GTTGCCAGTTGCCGAACACCGGCA-3′<br>5′-AAACTGCCGGTGTTCGGCAACTGG-3′ |
| Genotyping of *OsC4H1-KO* and *OsC4H1/2a/2/b-TKO* | 5′-CTATCGAGACGACGCTGTGG-3′<br>5′-TGGTGTCCACCTTGTCCATC-3′ |
| Genotyping of *OsC4H2a/2b-DKO* and *OsC4H1/2a/2/b-TKO* | 5′-GTGAGCGGGTCTAGTTCGAG-3′<br>5′-CGACACCACCACCAGGTT-3′ |
| Genotyping of *OsC4H2a/2b-DKO KO* and *OsC4H1/2a/2/b-TKO* | 5′-AAACGACATCGGTCCCAGT-3′<br>5′-CGACACCACCACCAGGTT-3′ |
| Off-target analysis of *C4H1* guide 2 (site 1-1) | 5′-TACAGAACCCACCAGCAATG-3′<br>5′-CAGGTACACCATGCGAACAG-3′ |
| Off-target analysis of *C4H1* guide 2 (site 1-2) | 5′-GCGCTTTCTCTTGGAAACAC-3′<br>5′-CACTTGCACAATGGAGGATG-3′ |
| Off-target analysis of *C4H1* guide 2 (site 1-3) | 5′-GGAGTTGCGGTAGCTTCAGT-3′<br>5′-GCGACGACCATGAGAGTGTA-3′ |
| Off-target analysis of *C4H1* guide 6 (site 2-1) | 5′-TCGTGGAGAACATCAACGTG-3′<br>5′-CCTTAATCTGCTCCTGACACAA-3′ |
| Off-target analysis of *C4H1* guide 6 (site 2-2) | 5′-TCGTGGAGAACATCAACGTG-3′<br>5′-AATTGGATTTATCGACTGATGG-3′ |
| Off-target analysis of *C4H1* guide 6 (site 2-3) | 5′-GCGTCTGCAAGGGTATCTTC-3′<br>5′-GCCTCGCTCAAGTACTGCTC-3′ |
| Off-target analysis of *C4H2a/2b* guide 2 (site 3-1) | 5′-ATCACAGGAGCCTGACACG-3′<br>5′-GAGCTGTCCTGACCTCAACC-3′ |
| Off-target analysis of *C4H2a/2b* guide 2 (site 3-2) | 5′-CCGAGGAGACCATCGAGAG-3′<br>5′-TGAGAAGTCTGTCACCACAACA-3′ |
| Off-target analysis of *C4H2a/2b* guide 2 (site 3-3) | 5′-TTGGTCTCCATGATTTCCTC-3′<br>5′-AGCTGGAGTTGGAGATGGTG-3′ |
| Off-target analysis of *C4H2a/2b* guide 6 (site 4-1) | 5′-GTTGTGCCTTTCCTCAGGTC-3′<br>5′-GGACCTCCGAACACTAGATCC-3′ |
| Off-target analysis of *C4H2a/2b* guide 6 (site 4-2) | 5′-GCTACTGGCCACTTCGTCAT-3′<br>5′-CCTGCCCGACGAATATCTTA-3′ |
| Off-target analysis of *C4H2a/2b* guide 6 (site 4-3) | 5′-GCCCTCCACATTAGCCATAG-3′<br>5′-ACGTACCCTGACGAAGCAGT-3′ |
